# Antirrhinum flower shape: unravelling gene expression across developmental axes and boundaries

**DOI:** 10.1101/2025.11.10.687484

**Authors:** Ana Maria Cunha, João Raimundo, Alexandra Verweij, Desmond Bradley, Enrico Coen, Maria Manuela Ribeiro Costa

**Affiliations:** Centre of Molecular and Environmental Biology (CBMA), University of Minho, Campus de Gualtar, 4710-057 Braga, Portugal; Department of Cell and Developmental Biology, John Innes Centre, Colney Lane, Norwich NR4 7UH, UK; Católica Biomedical Research Centre (CBR), Universidade Católica Portuguesa, Lisbon, Portugal; Enza Zaden, Research and Development B.V., P.O. box 7, 1600 AA Enkhuizen, The Netherlands

**Keywords:** Snapdragon, Flower Development, Transcriptomics, Dorsoventral asymmetry, Transcription Regulation

## Abstract

- Zygomorphic flowers in *Antirrhinum majus* arise from coordinated patterning along the dorsoventral and proximal–distal axes, but the transcriptional programs that pattern these axes to generate the complex corolla morphology remain poorly understood.
- We performed comparative transcriptomic analyses of dorsal and ventral petals across developmental stages, combined with expression profiling in floral symmetry mutants, to identify genes potentially associated with dorsal identity (*AmCYC*-dependent) and ventral identity (*AmDIV*-dependent), and to map their spatial dynamics along the proximal–distal axis. Axis-specific and boundary-localized expression patterns were validated by in situ hybridisation.
- We identified dorsal-specific candidate targets of *AmCYC* and ventral-specific genes regulated by *AmDIV*, including factors with regionally restricted expression along the proximal–distal axis. We further found that a conserved *NGATHA-LIKE1–BRASSINAZOLE-RESISTANT 1–miR164* module, known for primordia growth regulation in *Arabidopsis*, has been co-opted in *Antirrhinum* to act as a potential spatiotemporal regulator of *AmCUP,* a target of AmDIV, thereby contributing to the complex shape of the *Antirrhinum* flower.
- Our results provide new insight into the gene regulatory landscape underlying zygomorphic flower development and highlight specific regulatory modules that may coordinate axis patterning with boundary establishment in the corolla. This work establishes a framework for further understanding how developmental gene networks shape a complex corolla morphology and offers new targets for functional and evolutionary investigation of floral diversity.

## Introduction

The establishment of asymmetrical patterns of gene expression is known to control the three-dimensional shape of biological structures, including flowers. In Snapdragon (*Antirrhinum majus*), the development of the corolla leads to a specific arrangement and form consisting of 5 petals, fused in a tube that then separate forming three types of petals shapes, dorsal, lateral and ventral (Fig. 1a). This flower requires the establishment of dorsoventral asymmetry along the floral meristem through the action of a specific set of genes (Luo et al., 1996; Luo et al., 1999; Galego and Almeida, 2002; Corley et al., 2005; Costa et al., 2005). Genes expressed asymmetrically along the dorsoventral axis of the flower act in combination with genes along the proximal-distal axis to form the complex corolla shape (Green et al., 2010; Rebocho et al., 2017a; Rebocho et al., 2017b). However, the knowledge of the different transcriptional activities along the dorsoventral axis of the flower, and how different spatiotemporal patterns of gene expression modify cellular outcomes along the proximal-distal axis of the petals remains limited. Here we apply a transcriptomic approach, coupled with *in situ* validation, to identify additional genes regulating the axes of development, which ultimately shape the asymmetric *Antirrhinum* corolla.

**Figure 1.**
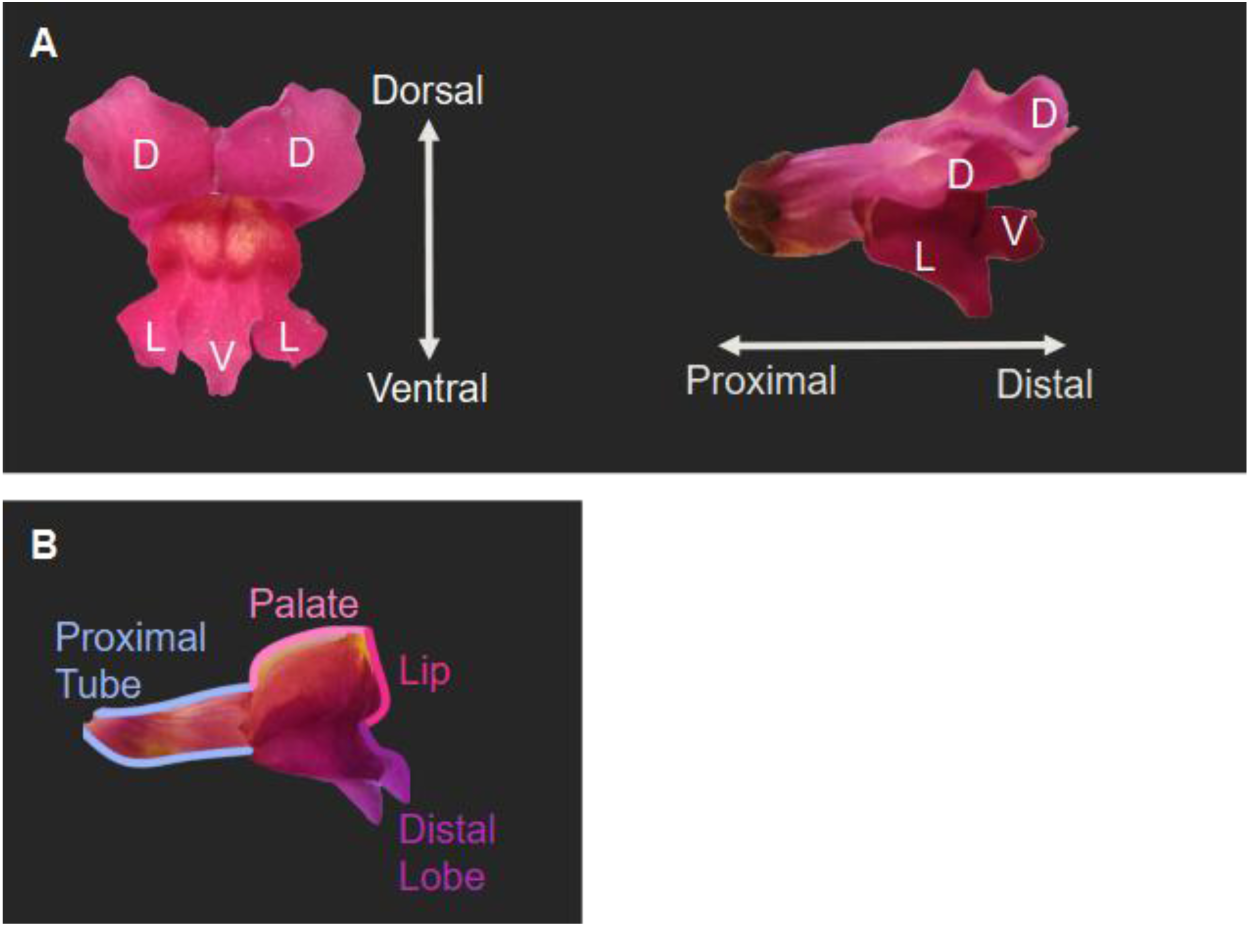
*Antirrhinum* flower shape. **(A)** The dorsoventrally asymmetric corolla consists of two large dorsal petals, two smaller lateral petals, and a single small ventral petal. The proximal regions of the petals are fused, forming a tube. **(B)** The lower corolla is composed by the ventral and lateral petals and has a complex shape along the proximal-distal axis. This shape is characterised by its wedge-like fold created by the palate and lip.

The establishment of flower dorsoventral asymmetry in *Antirrhinum* is coordinated by a set of transcription factors including *CYCLOIDEA (AmCYC)*, *DICHOTOMA* (*AmDICH*), *RADIALIS* (*AmRAD*) and *DIVARICATA* (*AmDIV*) (Luo et al., 1996; Almeida et al., 1997; Luo et al., 1999; Galego and Almeida, 2002; Corley et al., 2005; Costa et al., 2005). *AmCYC* is expressed in the dorsal regions of the flower meristem (FM), starting from stage one of development onwards (Luo et al., 1996; Clark and Coen, 2002). In early stages of flower development, *AmCYC* sets dorsal identity by retarding the growth of dorsal regions of the FM (Luo et al., 1996). Later, the role *of AmCYC* is whorl specific, promoting the growth of the dorsal petals and arresting the development of the dorsal stamen (Luo et al., 1996). *AmCYC* has a paralogue, *AmDICH,* that also controls the development of the dorsal petals specifically the shape of their dorsal half (Luo et al., 1999). Another dorsal gene *AmRAD*, is a downstream target of AmCYC, expressed in the same domain as *AmCYC* and expressed soon after *AmCYC* (Corley et al., 2005). The *cyc* mutant displays a strong phenotype: the dorsal petals assume some lateral characteristics and the rest of the petals have ventral identity (Carpenter and Coen, 1990; Luo et al., 1996). In *dich* mutants there is a loss of internal asymmetry of the dorsal petal (Luo et al., 1999). In the double mutant *cyc dich* dorsal identity is completely lost, and the flowers become radially symmetric, with all the petals resembling the wild-type ventral petal (Luo et al., 1999). Similarly, *rad cyc* double mutants are fully ventralised, however, the strongest loss-of-function *rad* mutant retains some dorsal identity, suggesting that *AmCYC* regulates additional targets other than AmRAD (Corley et al., 2005). In early flower development, *AmDIV* is expressed across the FM and its activity increases the length of lateral and ventral petals (Galego and Almeida, 2002). Later, *AmDIV* is asymmetrically expressed in the ventral furrow, with higher transcript levels in the adaxial epidermis than in the other layers. The formation of this fold depends on *AmDIV* influencing tissue growth parallel to the proximal-distal axis in the primordial palate and lip regions, giving the lower corolla its characteristic complex shape (Green et al., 2010; Rebocho et al., 2017a; Rebocho et al., 2017b). *AmDIV* is also responsible for the initiation of distinct cell types, including the development of trichomes (Perez-Rodriguez et al., 2005). In semidominant *div* mutants, the ventral petal adopts a lateral identity and *cyc dich div* mutants result in radially symmetric flowers with an all-lateral petals phenotype, which demonstrates the role of *AmDIV* in determining ventral identity (Almeida et al., 1997). Loss-of-function *div* mutants also fail to form the wedge-shaped fold of the ventral petal creating a less complex petal shape (Almeida et al., 1997; Galego and Almeida, 2002). *AmDIV* only affects the growth of ventral and lateral petals, despite its expression in all petals (Galego and Almeida, 2002). In dorsal petals, the presence of *AmRAD* disrupts the interaction of AmDIV with the protein DIV-and-RAD-Interacting-Factor (AmDRIF), which in turn prevents AmDIV action (Raimundo et al., 2013). The regulatory network of genes controlled by *AmDIV* during petal development remains poorly understood despite its broad functions. The boundary domain gene *CUPULIFORMIS* (*AmCUP*) has been identified as an AmDIV direct target during ventral petal development (Rebocho et al., 2017a). Flowers of *cup* mutants lack a complex shape, without a visible palate or lip, and exhibit inter-whorl fusions (Weir et al., 2004; Rebocho et al., 2017a). *AmDIV* promotes the expression of *AmCUP* in the ventral furrow, contributing to tissue growth and the formation of the palate and lip region (Rebocho et al., 2017a). These growth-enhancing effects of *AmCUP* may involve the modulation of auxin through *AmYUCCA1*, an auxin biosynthesis gene. After petal initiation, *AmCUP* expression represses growth in two domains: a proximal domain that creates the inter-whorl boundary, and a distal domain that separates the petals. The role of *AmCUP* in floral organ boundary domains seems independent of *AmDIV* expression suggesting that other genes might be involved in *AmCUP* regulation. *AmCUP* is a member of the NAC-domain family of genes (Weir et al., 2004) that contains other members recognised for their roles in defining organ boundaries during flower development in various species such as *Arabidopsis* (Aida et al., 1997; Hibara et al., 2006), petunia (Souer et al., 1996; Zhong et al., 2016), tomato (Berger et al., 2009; Hendelman et al., 2013) and *Medicago* (Cheng et al., 2012). NAC-domain genes have been extensively researched in leaf organogenesis, where they perform a dual function, repressing growth within their expression domain, and promoting growth outside of these domains by modulating auxin flux (Blein et al., 2008; Kawamura et al., 2010; Bilsborough et al., 2011; Hasson et al., 2011; Bhatia et al., 2023). In *Antirrhinum* and other species with dorsoventral asymmetry, *Linaria* and *Mimulus*, *AmCUP* appears to have been recruited for the modulation of the complex shape of the corolla, under the control of dorsoventral genes (Rebocho et al., 2017a).

The developmental programme that establishes dorsoventral asymmetry has been recruited numerous times and independently in other species to control flower development (Cubas, 2004; Busch and Zachgo, 2009; Preston and Hileman, 2009; Rosin and Kramer, 2009; Preston et al., 2011; Hileman, 2014; Wessinger and Hileman, 2020). Nevertheless, the understanding of the genetic network regulating dorsoventral asymmetry remains limited. Transcriptome analyses, aiming to explore dorsoventral asymmetry in petals have so far been focused to species with less complex-shaped corollas, such as *Sinningia speciosa* and *Pisum sativum* (Jiao et al., 2017; Pan et al., 2022) and have focused on the role of *CYC* homologues. In *Antirrhi*num, a recent transcriptomic study examined only small portion of the ventral petal, a curvature of the flower tube named the gibba, in comparison with the development of a similar structure in *Linaria vulgaris,* the spur (Cullen et al., 2023).

Here we employ a multi-level transcriptome analysis approach to expand the known gene network responsible for flower development in *Antirrhi*num. The present study identified genes that exhibit differential expression along the dorsoventral axis in an early stage of the FM stage and during petal development. Additionally, corolla transcriptomes of flower symmetry mutants were analysed in order to identify AmDIV targets. Promoter analysis on newly identified genes revealed putative direct targets for AmCYC in the dorsal meristem and for AmDIV in the ventral petals. Transcriptomes of different parts of the ventral petal along its proximal-distal axis, including the palate and lip, provided information of gene expression for later stages of corolla development along this axis. Some of these genes were selected for further analysis by *in situ* hybridisation revealing their specific expression in the adaxial epidermis of the ventral petal. Genes identified in the transcriptomic analysis as possible regulators of *AmCUP* were also studied by *in situ* hybridisation during flower development. This revealed the existence of a potentially conserved regulatory module that controls organ boundaries in other species and was co-opted to control petal development in *Antirrhinum*.

## Materials and Methods

### Plant materials and growth conditions

Snapdragon (*Antirrhinum majus*) wild-type plants (JIC Stock 7) and the mutants *cyc-608 dich719* (JI718), *cyc-608 dich^G^ div-80* (JI348) (Green et al., 2010) and *cup^sem^* (Weir et al., 2004) were grown in a greenhouse at the John Innes Centre. The development stage of the bud was determined through size and morphology (Vincent and Coen, 2004; Rebocho et al., 2017a). For the transcriptome analysis along the dorsoventral axis, the following tissues were collected in triplicate: dorsal, lateral, and ventral petals dissected from a 5 mm-wide wild-type bud; whole corollas of the radially symmetric mutants *cyc dich* and *cyc dich div,* from 5 mm-wide buds (Supplementary Table 1). To study an earlier stage of development, dorsal and ventral regions of a flower meristem at 5 days after floral meristem initiation (DAI) were sampled in duplicate using laser microdissection. Similarly, four ventral regions (tube, palate, lip, lobe) of *cyc dich* were collected in duplicate using laser microdissection in order to study gene expression along the proximal-distal axis. For the analysis of *cup* mutants, 3.5 mm-wide buds of *cup* and wild-type plants were sampled in duplicate. Every sample was immediately frozen in liquid nitrogen and ground to a fine powder. RNA was extracted using the RNeasy Kit (QIAGEN) according to the manufactureŕs instructions.

### Analysis of transcriptomic data

RNA samples were sent for sequencing varying in read size and library type (Supplementary Table 1). The Galaxy server (usegalaxy.org) was used for data processing and analysis. Quality control analysis was performed using the FastQC software (Andrews, 2023) 0.12.1 followed by TrimmGalore! (Krueger, 2023) V0.6.7 for read trimming and adaptor removal. The reads were then mapped to the Antirrhinum genome Version 3 (Li et al., 2019) using HISAT2 (Kim et al., 2015) version 2.1.1. Mapped reads were quantified using htseq-counts (Anders et al., 2015) version 0.9.1. The assessment of differential gene expression between samples was conducted using DESeq2 (Love et al., 2014) version 2.11.40.8. Genes were considered as differentially expressed if false discovery rate (FDR) < 0.01 and | log2(fold change) | > 1.5. To better understand the roles of the identified genes, their corresponding protein sequences were used in a basic local alignment search tool (BLAST) (Altschul et al., 1990) against the database UniProtKB/Swiss-Prot with the taxonomy filter of Viridiplantae (33090) and e-Value 1.0E-3. Additionally, Gene Ontology (GO) mapping and GO annotation were performed (Ashburner et al., 2000; Götz et al., 2008; Bateman et al., 2023).

### *In situ* hybridisation

RNA *in situ* hybridisation was performed according to Jackson (1992) with some modifications described in Coen et al. (1990) and Bradley et al. (1993). To generate specific digoxigenin-labelled riboprobes probes, the open reading frame of the genes *AmBZR1* (*Am01g00480*), *AmNGAL1* (*Am07g28680*) and *AmCUP* (*Am03g10700*) preceded by the sequence (5’-GTTGTAAAACGACGGCCAGTGAATTGTAATACGACTCACTATAGGGCGAATTGGG CCCGA-3’), having the M13F primer site and the T7 promoter, were synthesised by Strings DNA Fragments (Invitrogen). Five hundred nanograms of the synthesis reaction were labelled using T7 RNA polymerase (10881767001, Roche) and DIG-UTP (11209256910, Roche), according to the instructions of the manufacturer. The *miR164* RNA probe (TGCACGTGCCCTGCTTCTCCA) was ordered from Exiqon, and the *in situ* hybridisation protocol optimised according to Javelle and Timmermans (2012). The DIG-labelled probes were hybridised to tissue sections of inflorescences from *Antirrhinum* wild type. To visualise the DIG-labelled probes an immunochemical assay was performed using an anti-DIG antibody that was conjugated with alkaline phosphatase.

### *In silico* analysis of promoter regions for binding sites

Promoters of genes more expressed in the dorsal region of the flower meristem were scanned for AmCYC binding sites. Similarly, promoters of genes more expressed in the ventral petals were scanned for AmDIV binding sites. Promoter sequences were defined as 1000 bp before the initiation codon. For the identification of the binding sites, promoters were analysed using the Find Individual Motif Occurrences (FIMO v5.5.5) tool (Grant et al., 2011) from the MEME suite (Bailey et al., 2015) online (meme-suite.org/meme/tools/meme). The motifs previously identified for AmCYC binding 5’-GGNCCCNC-3’ (Costa et al., 2005) and AmDIV binding 5’-GATAAG-3’ (Raimundo et al., 2013) were supplied as input in FIMO, and only exact matches were considered. Motif enrichment analysis of CYC binding sites was conducted by comparing 29 CYC target promoters (positive sequences) with 25 DIV target promoters (negative sequences) using the Analysis of Motif Enrichment (AME) tool from the MEME suite. The *AmCUP* promoter was scanned for the previously determined binding site of the BZR1 homologue of Arabidopsis thaliana 5’-CGTGCG-3’ (He et al., 2005).

## Results

### Dorsal-ventral differentially expressed genes in two stages of flower development

The development of the *Antirrhinum* flower requires the asymmetric expression of genes along its dorsoventral axis. To expand our knowledge of the genetic network that controls flower dorsoventral asymmetry, a transcriptomic analysis was performed on dorsal and ventral tissues. For an early stage of development, a FM was identified at 5 DAI (days after floral meristem initiation) according to Rebocho et al. (2017a). At this stage sepal primordia in the FM were evident as described in Vincent and Coen (2004). The dorsal and ventral regions of the FM were separately isolated by laser microdissection (Fig. 2a) and used for transcriptomic analysis. The transcriptomes of dorsal and ventral regions of the FM were compared and genes with a False Discovery Rate (FDR)<0.01 and |log2(fold change)|>1.5 were considered as differentially expressed genes (DEGs).

**Figure 2.**
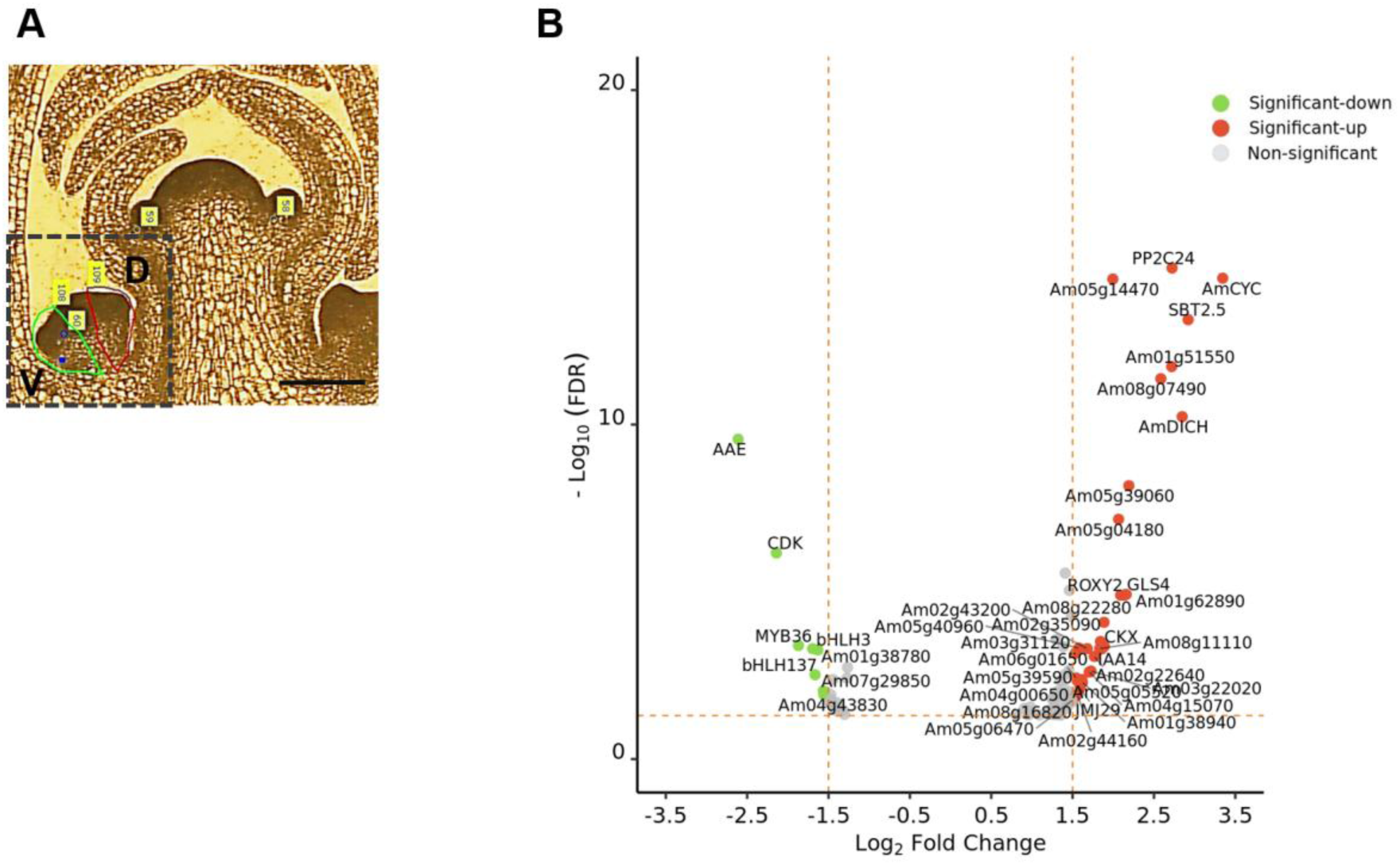
Transcriptome analysis of dorsal and ventral regions from an *Antirrhinum* flower meristem. **(A)** From a wax embedded section of an inflorescence, a flower meristem at five days after floral meristem initiation (DAI; dashed box) was laser micro dissected into dorsal (D; red) and ventral (V; green) regions and used for transcriptomic analysis. Scale 150 µm. **(B)** Volcano plot with differentially expressed genes (DEGs) between dorsal and ventral regions of the FM. Genes in colour were considered significantly differential expressed if FDR<0.01 and |log2(fold change) |>1.5.

A total of 29 genes were identified as more expressed in the dorsal region of the FM, while 7 were more expressed in the ventral region (Fig. 2b; Supplementary Dataset 1.1). Among the genes more expressed dorsally, were *AmCYC* and *AmDICH*, which are known dorsal-specific genes (Luo et al., 1996; Luo et al., 1999), demonstrating the specificity and good quality of the microdissection of the FM regions. Other more dorsally expressed genes included a *MADs-box* gene identified in *Antirrhinum* as *AmDEFH7* (Li, 2002; Coenen et al., 2018) and a homologue of the glutaredoxin *ROXY2* from *Arabidopsis thaliana*. The expression also of homologues of *CYTOKININ DEHYDROGENASE* (*CKX*) and *INDOLEACETIC ACID-INDUCED 14* (*IAA14*) suggested the involvement of cytokine catabolism and auxin response in the development of the dorsal region. A gene related to cell wall synthesis, callose synthase *GLUCAN SYNTHASE-LIKE 4* (*GSL4*), and an epigenetic regulator *JUMONJI DOMAIN-CONTAINING 29* (*JMJ29*), were also identified as more dorsally expressed. At this stage of development, seven genes were more expressed in the ventral region of the FM. These included an *ACETYLAJMALAN ESTERASE* (*AAE*), a *CALCIUM-DEPENDENT KINASE* (*CDK*), and homologues of members of the *MYB* and basic helix-loop-helix (*bHLH*) family of transcription factors, *MYB36*, *bHLH137* and *bHLH3*.

*AmCYC* has a crucial role in setting dorsal identity from early stages of flower development (Luo et al., 1996). Thus, to evaluate if some of the dorsally expressed genes could be direct targets of *AmCYC,* their promoter sequences were analysed. Sequences of 1 kb before the ATG codon were selected and scanned for AmCYC binding sites (GGNCCCNC) as defined by Costa et al. (2005). Of a total of 29 promoters scanned, 11 had AmCYC binding sites (Supplementary Table 2), including one in the *AmCYC* promoter that was previously described by Costa et al. (2005). Motif enrichment analysis has confirmed that CYC binding sites are significantly enriched (*p*-value=3.16^e-2^) in promoters of CYC target genes by comparing to those of putative DIV targets (dotted box in Fig. 4c). Other genes with AmCYC binding sites in their promoters included those of glutaredoxin *ROXY2* and callose synthase *GSL4*.

To identify dorsal, ventral and lateral petal specific genes at a later stage of flower development, 5 mm-width buds were used to extract RNA from the different petal types (Fig. 3a). When comparing the transcriptomes of dorsal and ventral petals, 37 genes were more expressed in dorsal petals, whereas 68 were more expressed in ventral petals (Fig. 3b; Supplementary Dataset 1.2). A comparison between dorsal and lateral petals yielded 21 DEGs (Supplementary Dataset 1.3), 20 of which were common to the dorsal-ventral comparison. In contrast, no DEGs were found when comparing ventral and lateral petals. This could be because the dissection between lateral and ventral petals was done at the latero-ventral junction, where some genes responsible for ventral identity such as *AmDIV* and *AmCUP* are expressed (Rebocho et al., 2017a; Rebocho et al., 2017b).

**Figure 3.**
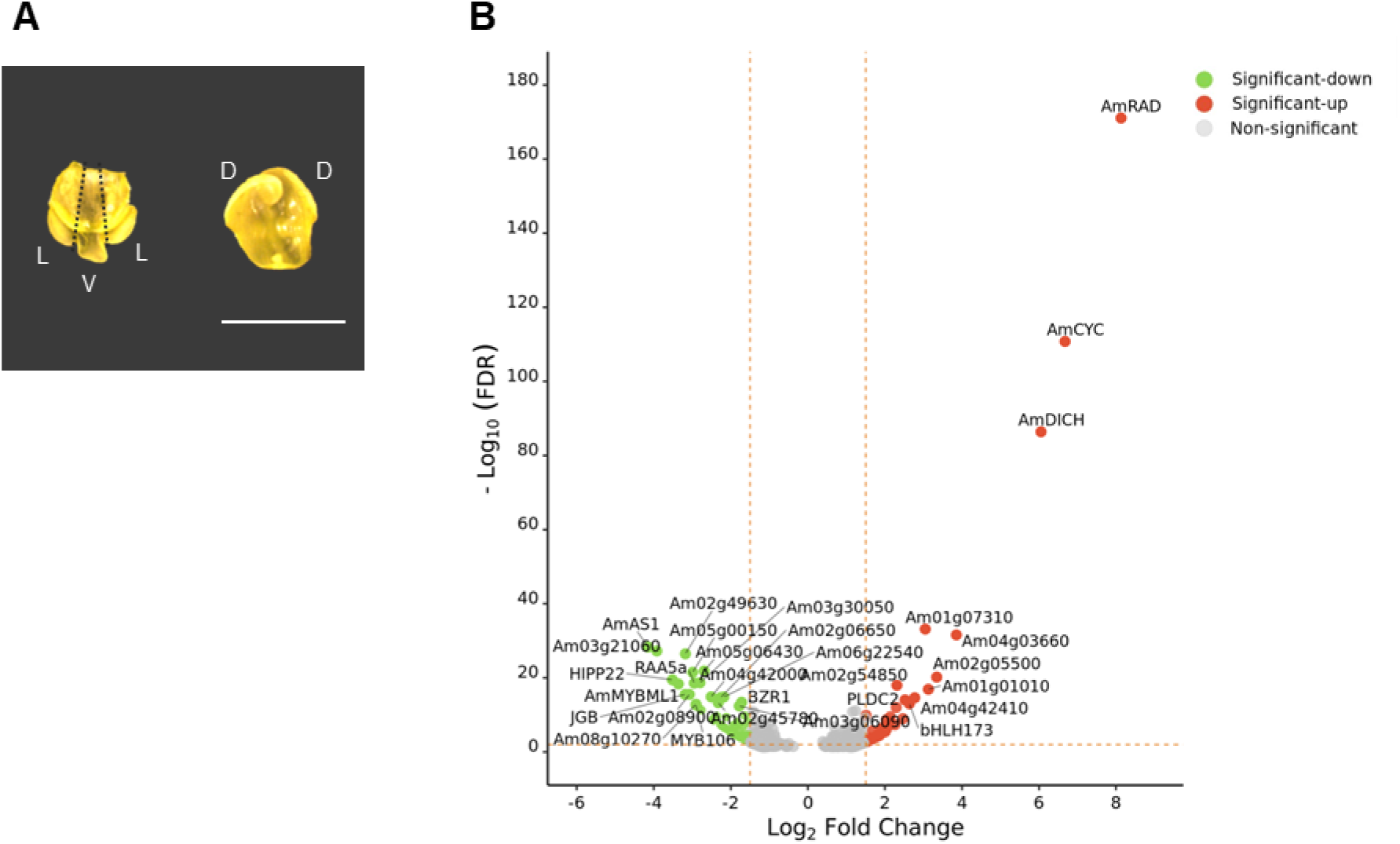
Transcriptome analysis of dorsal and ventral petals from a young *Antirrhinum* flower bud. **(A)** Dorsal (D), lateral (L) and ventral petals (V) were dissected from a pool of 5 mm-wide buds and used for transcriptomic analysis. Scale 5 mm **(B)** Volcano plot with DEGs between dorsal and ventral petals. Genes in colour were considered significantly differential expressed if FDR<0.01 and |log2(fold change)|>1.5.

In the petal samples, *AmCYC*, *AmDICH* and *AmRAD* are the most significantly DEGs being 103, 66 and 280 times more expressed in dorsal petals, respectively. *AmCYC* and *AmDICH* were the only genes in common between both analysis (FM and petals) using the cut-offs FDR<0.01 and |log2(fold change)|>1.5 (Supplementary Figure 1). *AmRAD* was significantly differentially expressed between dorsal and ventral petals of buds but not between the dorsal and ventral regions of the FM, suggesting a later up-regulation. Three genes more expressed in ventral petals were also previously described in the literature. The expression of *AmCUP*, *MYB MIXTA LIKE 1* (*AmMYBML1*) and *AUREUSIDIN SYNTHASE* (*AmAS1*) are crucial for the ventral petal development in terms of shape (Rebocho et al., 2017), cell types (Perez-Rodriguez et al., 2005) and colour (Davies et al., 2006), respectively. It is worth noting that another gene, *Am02g05500,* was identified as a possible homologue of *AmAS1,* however, it was more dorsally expressed. The identification of *AmCYC*, *AmDICH* and *AmRAD* as more expressed in the dorsal petals, and *AmCUP*, *AmMYBML1* and *AmAS1* in the ventral petals supports the methodology and quality of the data set.

GO analysis was applied to the remaining identified dorsal and ventral genes to better understand their function (Supplementary Dataset 1.4). Genes annotated with GO Terms related to flower development included homologues of: *NGATHA-LIKE 1 (NGAL1), ENHANCER-OF-JOINTLESS-2 (EJ2)* and *PHOSPHOINOSITIDE PHOSPHOLIPASE C 2 (PLCD2)*, more expressed in dorsal petals; and *CEN-like protein 1* (*CENL1)*, *SUPERMAN (SUP)*, *GATA9* and *LOB DOMAIN-CONTAINING PROTEIN 36 (LBD36)*, more expressed in ventral petals. Since dorsoventral asymmetry in *Antirrhinum* may also be influenced by hormones (Bergbusch, 1999; Rebocho et al., 2017a), a search was performed to identify genes with GO terms associated with hormones. This identified homologues of genes involved in the biosynthesis of cytokinin *LONELY GUY 5* (*LOG5*), brassinosteroids signalling *BRASSINAZOLE-RESISTANT 1* (*BZR1*), *PACLOBUTRAZOL RESISTANCE 6* (*PRE6*)*, B-BOX ZINC FINGER PROTEIN 21* (*BBX21), bHLH173* and *NGAL1*, and auxin biosynthesis *PLCD2*.

In addition to *AmMYBML1*, homologues of *MYB16* and *MYB106* were also annotated as involved in trichome branching. In a phylogenetic analysis (Supplementary Fig. 2), MYB106 clustered with AmMYBL1, whereas MYB16 was phylogenetically more distant from the MYBML proteins*. MYB106* and *AmMYBML1* are likely the result of a recent duplication

### Differentially expressed genes between fully lateralised and ventralised mutants

In the *cyc dich* double mutant, ventral identity expands to all petals forming a radially symmetric fully ventralised flower (Luo et al., 1996; Luo et al., 1999) whereas in *cyc dich div,* the radially symmetric flower has all petals lateralised (Almeida et al., 1997) (Fig. 4a). To further corroborate the DEGs obtained from the comparison between dorsal and ventral petals, the transcriptome of the *cyc dich* petals was compared to dorsal wild-type (WT) petals. Genes more expressed in the ventral petal correlated with genes more expressed in the ventralised *cyc dich* mutant (Fig. 4b). The comparison between the transcriptomes of *cyc dich div* and *cyc dich* mutants returned a total of 1819 DEGs (Supplementary Dataset 1.5). The putative targets of *AmDIV* could be among genes more expressed in ventral petals and in *cyc dich* mutant petals, and downregulated in *cyc dich div* mutants. A total of 26 genes fulfilled these criteria and were considered as putative AmDIV targets (Fig. 4c). To distinguish which genes could be potential direct targets of AmDIV, the promoter region corresponding to the 1 kb before the initiation codon was analysed for AmDIV binding sites. AmDIV binds to I-box motifs, in particular to the consensus sequence 5’-GATAAG-3’ (Raimundo et al., 2013). Of the 26 ventral genes identified as potentially regulated by AmDIV, 13 contained AmDIV binding sites in their promoters (Fig. 4c; Supplementary Table 3). Promoters that contained more than one AmDIV binding sites included the ones of homologues of *BBX21, bHLH126, MYB16* and *HEAVY METAL-ASSOCIATED ISOPRENYLATED PLANT PROTEIN 22* (*HIPP22*). Other genes possibly regulated directly by AmDIV included homologues of *CARBONIC ANHYDRASE* (*CA*), *GATA9, ANTHOCYANIN 5-O-GLUCOSIDE-4’’’-O-MALONYLTRANSFERASE 2* (*5MAT2*), *ACETYLAJMALAN ESTERASE (AAE), RADL6 and JINGUBANG* (*JGB*).

**Figure 4.**
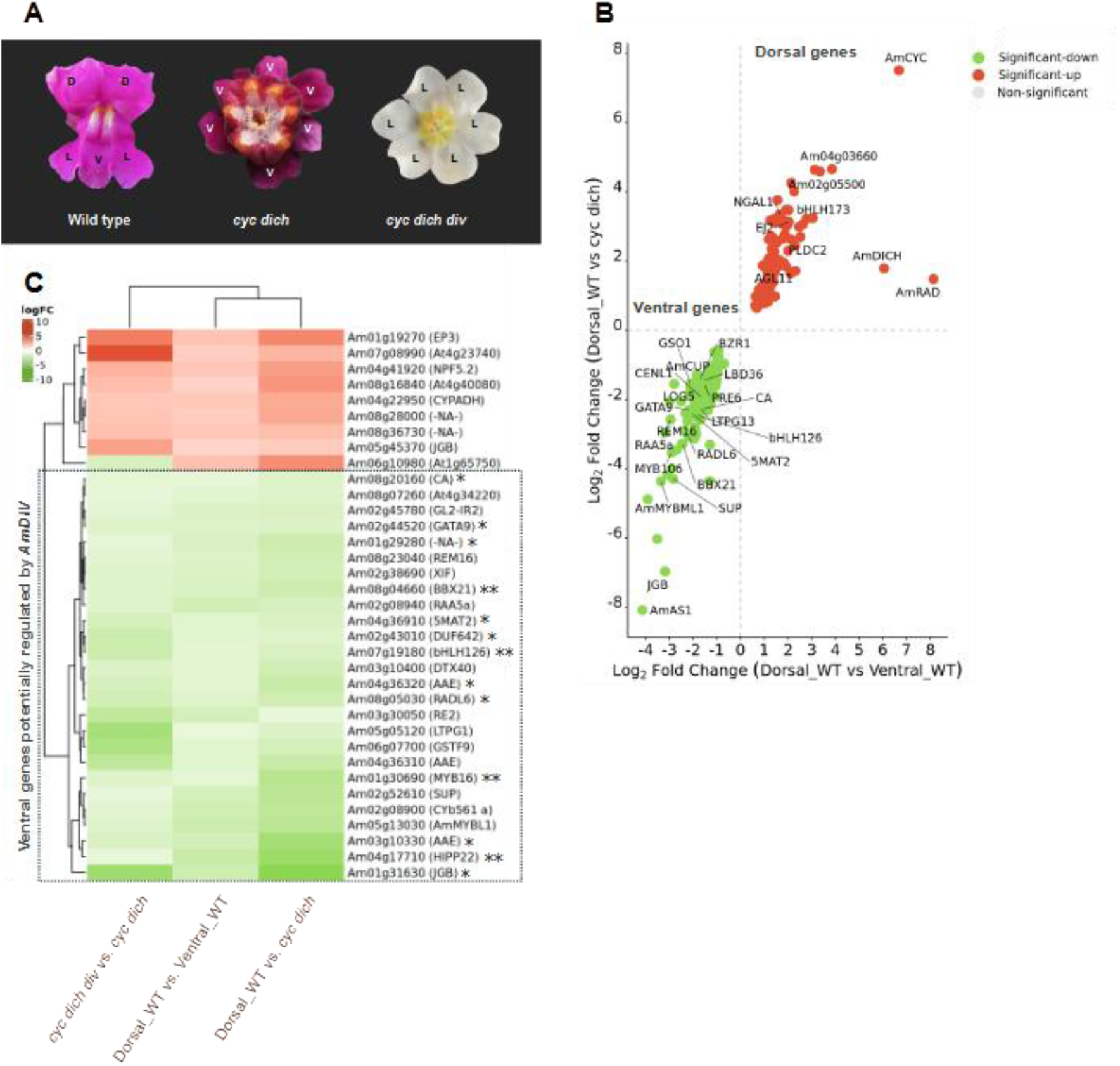
Transcriptome analysis of mutants with symmetrical flowers. **(A)** Wild-type (WT) *Antirrhinum* flowers have two dorsal (D) petals, two lateral (L) and one ventral (V) petal whereas the mutant *cyc dich* has six ventral petals, and *cyc dich div* six lateral petals. The pigmentation differences are independent effects of colour genes. **(B)** A correlation graph of DEGs common between WT dorsal vs ventral petals (x-axis) and WT dorsal petal with *cyc dich* mutant (y-axis). Genes coloured green correspond to genes more expressed in the ventral petal and the *cyc dich* mutant, while genes in red identify genes more expressed in the dorsal petal. Genes in grey do not fill the criteria of FDR<0.01 and are considered non-significant. **(C)** Heatmap of genes common to the three comparisons: WT petals dorsal vs. ventral, WT dorsal vs. *cyc dich* and *cyc dich div* vs. *cyc dich* (in this comparison, genes more expressed in the *cyc dich* mutant appear in green, whereas genes more expressed in the *cyc dich div* mutant appear in red). Genes possibly regulated by AmDIV are indicated inside the dotted box. Genes with one (*) or two (**) AmDIV binding sites in their promoters.

### Differentially expressed genes along the complex shaped ventral petal

The *Antirrhinum* ventral petal has a complex shape that relies on differential growth along the proximal-distal axis (Green et al., 2010; Rebocho et al., 2017a; Rebocho et al., 2017b). The proximal-distal axis can be divided into four regions with distinctive morphological features: the proximal tube, palate, lip and distal lobe (Fig. 1b). For simplicity, the proximal tube will be referred only as tube and the distal lobe only as lobe. Following the identification of DEGs across the dorsoventral axis, the goal was to determine if genes more expressed in the ventral petals were also differentially expressed across the proximal-distal axis. The petals of 6.5 mm-wide buds of the fully ventralised *cyc dich* mutant were dissected into tube, palate, lip, and lobe regions and their transcriptomes sequenced. The selection of this mutant aimed to maximize tissue acquisition to isolate the distinct parts of the ventral petal. The expression of the 68 genes previously identified as more expressed in the ventral petal than dorsal (Supplementary Dataset 1.2) were analysed across tube, palate, lip, and lobe regions. The normalised expression of genes across the samples were clustered and their patterns of expression visualised as a heatmap (Fig. 5a). Five clusters were identified based on the expression patterns, which correlated with expression in specific petal regions. The cluster of genes more expressed in lip-palate regions contained *AmCUP* and homologues of *GATA9, LBD36,* and *GASSHO1 (GSO1)*. As a control for clustering, 50 genes were randomly selected from the pool of genes that were expressed in all samples, and their expression was not associated with specific regions (Supplementary Fig. 3).

**Figure 5.**
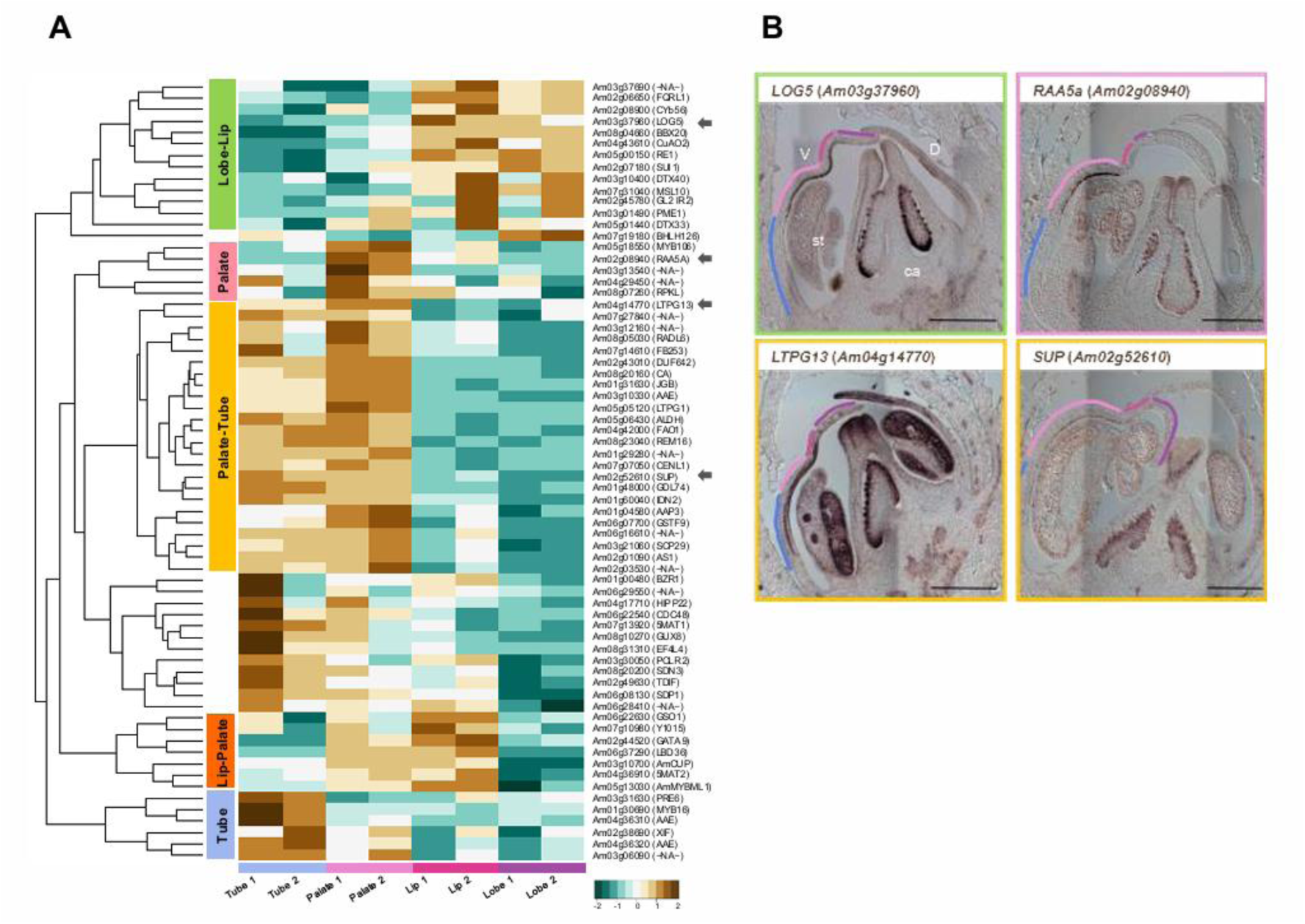
Transcriptomic analysis of micro-dissected ventral petal regions of the *Antirrhinum* flower. **(A)** Heatmap with hierarchical clustering of genes identified as more expressed in the ventral petal (from dorsal vs. ventral petal comparison), based on normalised expression across the regions of the tube, palate, lip, and lobe. Normalised expression of two biological replicates is shown. Genes marked with arrows were selected for further validation. **(B)** Gene expression by *in situ* hybridisation for Antirrhinum homologues of *LOG5* (top, left), *RAA5a* (top, right), *LTPG13* (bottom, left) and *SUP* (bottom, right) in middle sections of *A. majus* flowers. Letters correspond to ventral petal (V), Dorsal petal (D), stamens (st) and carpels (ca). Scale 500 µM.

Genes were selected from three of these clusters for further study through *in situ* hybridisation (ISH) in longitudinal sections through the midline of the flower at 14 DAI. These included the homologue of *LOG5* from the cluster of genes more expressed in the lobe-lip, *RAS-RELATED PROTEIN RABA5a (RAA5a)* from the palate cluster, and *LIPID TRANSFER PROTEIN GPI-ANCHORED 13 (LTPG13)* and *SUP* from the palate-tube cluster (Fig. 5b). *LOG5* was restricted to the adaxial side of the petal in the lobe, lip and palate regions, whereas *RAA5a* expression was limited to the adaxial side of the palate. *LTPG13* was detected in the tube, palate and to some extent in the lip. *SUP* expression was restricted to the palate in the adaxial side. All genes showed additional expression in carpel and/or stamen tissues. To demonstrate the absence of stickiness in the sampled tissues, sections from different stages of flower development hybridised with a representative sense probe are shown (Supplementary Figure 4).

In the ventral petal, some regions are characterised by the presence of yellow pigmentation and a higher density of trichomes. The yellow pigment aurone biosynthesis gene *AmAS1* (Davies et al., 2006) was more expressed in the tube and palate regions whereas genes associated with anthocyanin modification *5MAT2* and *5MAT1* (Suzuki et al., 2004, 2001), displayed higher expression in the lip-palate region, and palate-tube respectively (Fig. 5a). Other genes implicated in the anthocyanin and aurone biosynthesis pathway in *Antirrhinum* (Coen et al., 1986; Sommer and Saedler, 1986; Beld et al., 1989; Martin et al., 1991; Davies et al., 2006; Ono et al., 2006; Schwinn et al., 2006; Albert et al., 2021; Ichimura et al., 2021) were also analysed across samples (Supplementary Figure 3). Overall, genes involved in anthocyanin production were more expressed in the lip and lobe regions, characterised by a more intense magenta colour whereas genes involved in aurone biosynthesis were more expressed in the palate-tube regions. Similarly, *AmMIXTA* and genes belonging to the *MYBML* (*MIXTA-like*) family, which play roles in cell specification and trichome production (Glover et al., 1998; Perez-Rodriguez et al., 2005; Baumann et al., 2007; Jaffé et al., 2007) were analysed across samples. The *AmMIXTA* gene was more expressed in the lobe-lip region whereas *AmMYBML1* was more expressed in the lip-palate, together with the homologue of *MYB106* (Fig. 5a; Supplementary Figure 5). The other two members of the *MYBML* family, *AmMYBML2* and *AmMYBML3*, were more expressed in the palate and lobe respectively.

### Differentially expressed genes on the simplified-shaped flower of *cup* mutants

In *cup* mutant flowers, the corolla has a simpler shape lacking a visible palate or lip, implicating *AmCUP* in specifying late ventral identity (Rebocho et al., 2017). In order to identify genes potentially involved in the development of the complex corolla shape, a transcriptome analysis was conducted on 3.5 mm-wide WT and *cup* flower buds. This stage corresponds to the deepening of the ventral furrow that creates the characteristic palate and lip regions (Vincent and Coen, 2004). A total of 530 DEGs were obtained with 190 genes being down regulated in the *cup* mutant and 340 genes up regulated (Fig. 6a; Supplementary Dataset 1.6). Analysis of the top 20 GO terms revealed an involvement of several hormone-related genes including the hormones auxin, ethylene, abscisic acid and jasmonic acid, with genes both downregulated, as well as upregulated, in the *cup* mutants (Fig. 6b).

**Figure 6.**
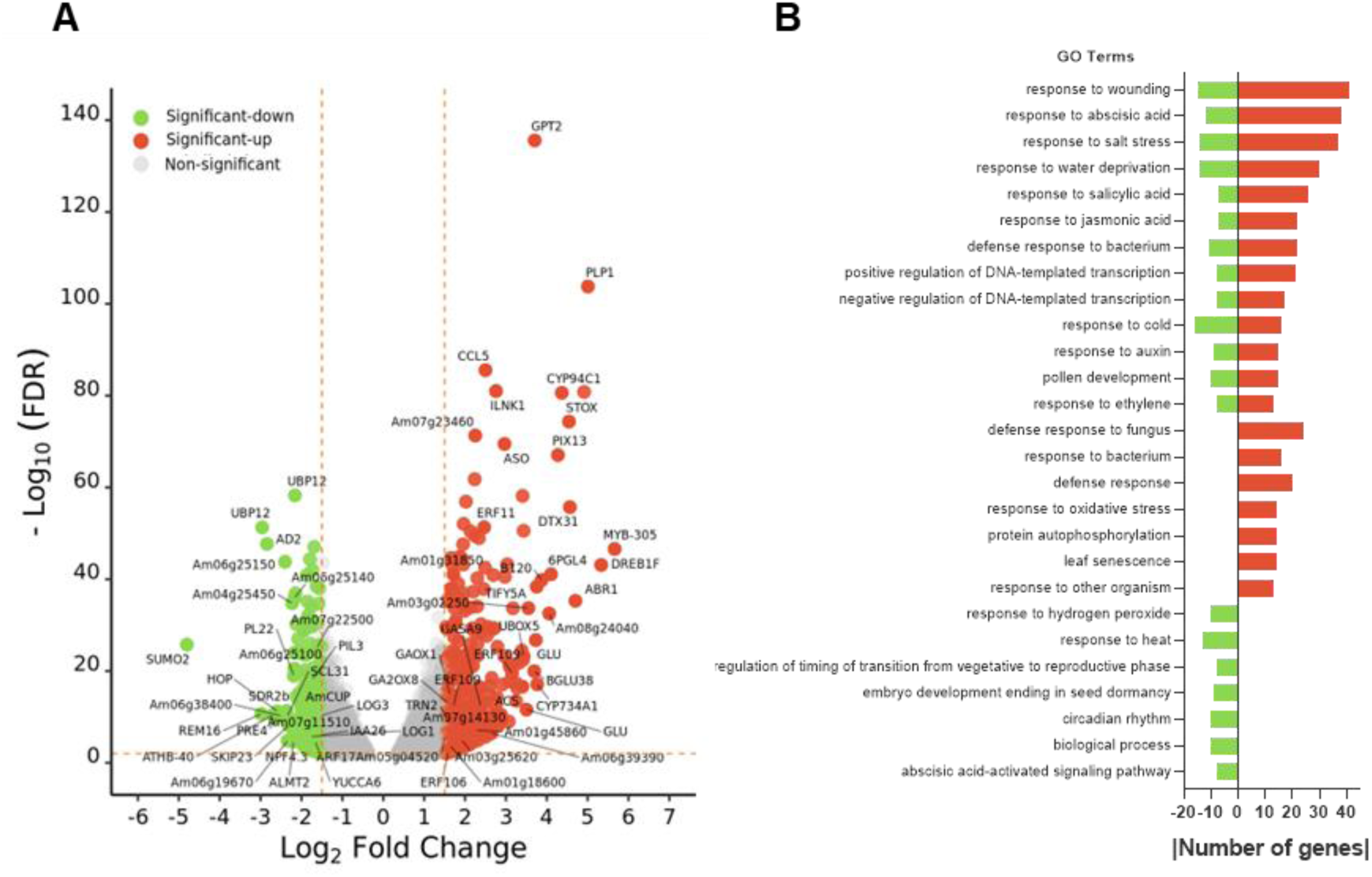
Transcriptomic analysis of *cup* mutant vs WT. **(A)** Volcano plot with DEGs between *cup* and WT buds 3.5 mm in width. Genes in colour were considered significantly differential expressed if FDR <0.01 and |log2(fold change) |> 1.5. **(B)** Top 20 Gene Ontology (GO) terms of genes downregulated (green) and upregulated (red) in the *cup* mutant. GO Terms shown belong to the Biological Process category and were ranked by the number of genes associated with each term.

Among downregulated genes were the homologues of the auxin biosynthesis gene *YUCCA6* and auxin transporter *PIN-LIKES 3* (*PIL3*) whereas another auxin transporter, *TORNADO 2* (*TRN2*), was upregulated. Similarly, the genes related to auxin signalling *AUXIN RESPONSE FACTOR 17* (*ARF17*) and *INDOLEACETIC ACID-INDUCED PROTEIN 26* (*IAA26*) were found to be downregulated, whereas *IAA4* was upregulated. DEGs associated with the biosynthesis of ethylene, gibberellin and cytokinin, were also identified in the *cup* vs WT comparison. Two homologues of ethylene biosynthesis genes, *1-AMINOCYCLOPROPANE-1-CARBOXYLATE SYNTHASE-LIKE* (*ACS*-*like*) were significantly up regulated in the mutant, as well as *GIBBERELLIN 2-BETA-DIOXYGENASE 1* (*GA2OX1*) and *GA2OX8,* involved in gibberellin catabolism and inactivation, respectively. Genes related to cytokinin biosynthesis, *LOG1* and *LOG3* homologues, were downregulated in the *cup* mutant. Some upregulated genes were assigned GO terms that relate to more than one hormone. For instance, *ETHYLENE RESPONSIVE FACTORS* (*ERFs*) transcription factors, were annotated as involved in ethylene, auxin and cytokinins, and a homologue of *GIBBERELLIN-REGULATED PROTEIN 9* (*GASA9*), was associated with gibberellin and brassinosteroids.

A gene downregulated in the *cup* mutant (Fig. 6a), homologue of *REPRODUCTIVE MERISTEM 16* (*REM16*), was also more expressed in the ventral petals and in the ventralised mutant *cyc dich* (Fig. 4c), and down regulated in *cyc dich div*. To elucidate tissue specificity and developmental timing of *AmREM16* expression, an ISH was performed at different stages of flower development. The expression of *AmREM16* at early stages of development, 6 and 7 DAI, localised in organ boundaries and at the basal part of the ventral sepal (Fig. 7a,b). At 12 DAI, *AmREM16* was restricted to the palate and the dorsal-dorsal junction (D-D) (Fig. 7d), while absent from the ventral-lateral (V-L) junction (Fig. 7c green bracket). *AmREM16* domains of expression were maintained at 14 DAI and it was also expressed later in ovule boundaries (Fig. 7e).

**Figure 7.**
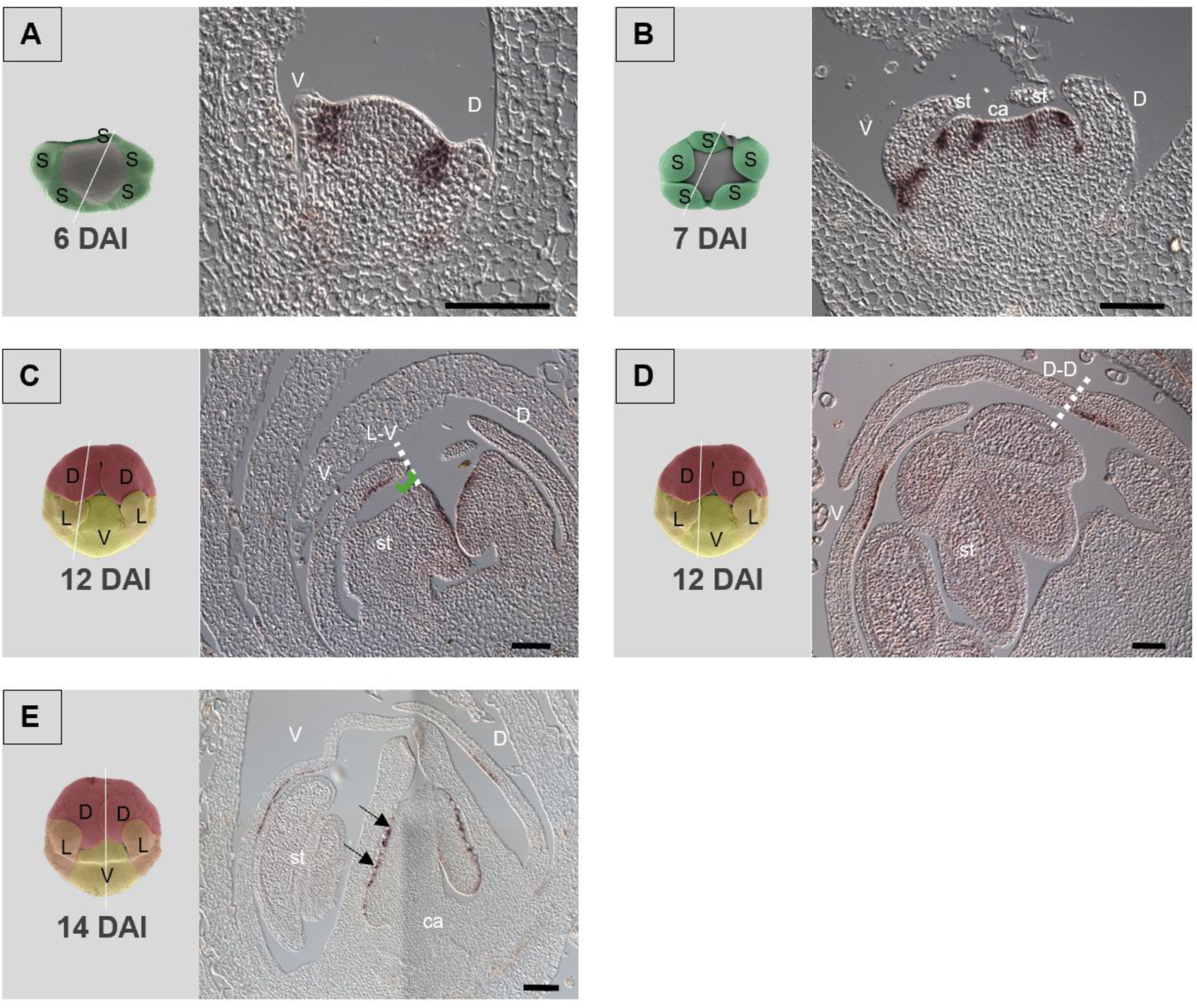
Expression of *AmREM16* during *Antirrhinum* flower development detected by *in situ* hybridisation. Sections of flower buds at different DAI were observed. **(A and B)** At 6-7 DAI, *AmREM16* was detected at organ boundaries, and in the base of the ventral sepal at 7 DAI. **(C and D)** At 12 DAI, *AmREM16* was expressed in the middle ventral petal and absent from the lateral-ventral junction (L-V dotted line; green bracket). *AmREM16* was observable in the dorsal-dorsal junction (D-D dotted line). **(E)** At 14 DAI, *AmREM16* was expressed in the palate region and in ovule boundaries (black arrows). D (dorsal); V (ventral); st (stamen); ca (carpel). Scale 100 µm.

### Expression of *miRNA164*, *AmNGAL1* and *AmBZR1* in comparison to *AmCUP*

In *Antirrhinum*, the only known regulator of *AmCUP* is AmDIV, which promotes the characteristic crescent-shaped expression of *AmCUP* in the ventral petal (Rebocho et al., 2017). In the comparison of dorsal and ventral petals, two genes were identified as homologues of *NGAL1* and *BZR1* (Fig. 3b), that were more expressed in the dorsal and ventral petals, respectively. In *Arabidopsis* these genes are known negative regulators of the expression of *AmCUP* homologues, the *CUCs* genes (Gendron et al., 2012a; Shao et al., 2020). Furthermore, *miRNA164* acts as a negative regulator of *CUCs* (Nikovics et al., 2006) to control leaf shape in *Arabidopsis* in a cross-species conserved module (Berger et al., 2009; Zheng et al., 2019; Wang et al., 2021a; Valoroso et al., 2022). To understand whether a similar network could regulate the expression of *AmCUP* in *Antirrhinum* during flower development, a side-by-side ISH was performed for *AmCUP* in comparison to *miRNA164*, *AmBZR1* and *AmNGAL1*.

The expression of *miR164* at 10 DAI, occurred at the distal tips of the developing petals, where the expression of *AmCUP* was not detected (Fig. 8a,b, purple brackets). At 12 DAI, *miR164* expression could be found at the distal tip of the ventral petal and across the dorsal petal. At the same time, the crescent-shaped expression of *AmCUP* was observed in the intermediary area of the ventral petal (Fig. 8b, blue bracket). By 14 DAI, this area of *AmCUP* expression extended further, giving rise to the palate and lip regions, which are separated by the rim and overlapped with *miRNA164* expression (Fig. 8c). At petal boundaries, *miR164* expression was visible in the V-L junction, whereas *AmCUP* expression was not detected (Fig. 8d, green bracket). In the dorsal petals, *AmCUP* was confined to the proximal region, adjacent to the D-D junction, while *miR164* was restricted to the distal zone (Fig. 8c,d). In the carpel, *miR164* was expressed in the ovules while *AmCUP* seemed to be restricted to the ovule boundaries (Fig. 8c, black arrows). Furthermore, *miR164* was expressed at the tip of the carpel (Fig. 8c) and in the stamens since early stages (Fig. 8a), and later in pollen sacs (Fig. 8c,d).

**Figure 8.**
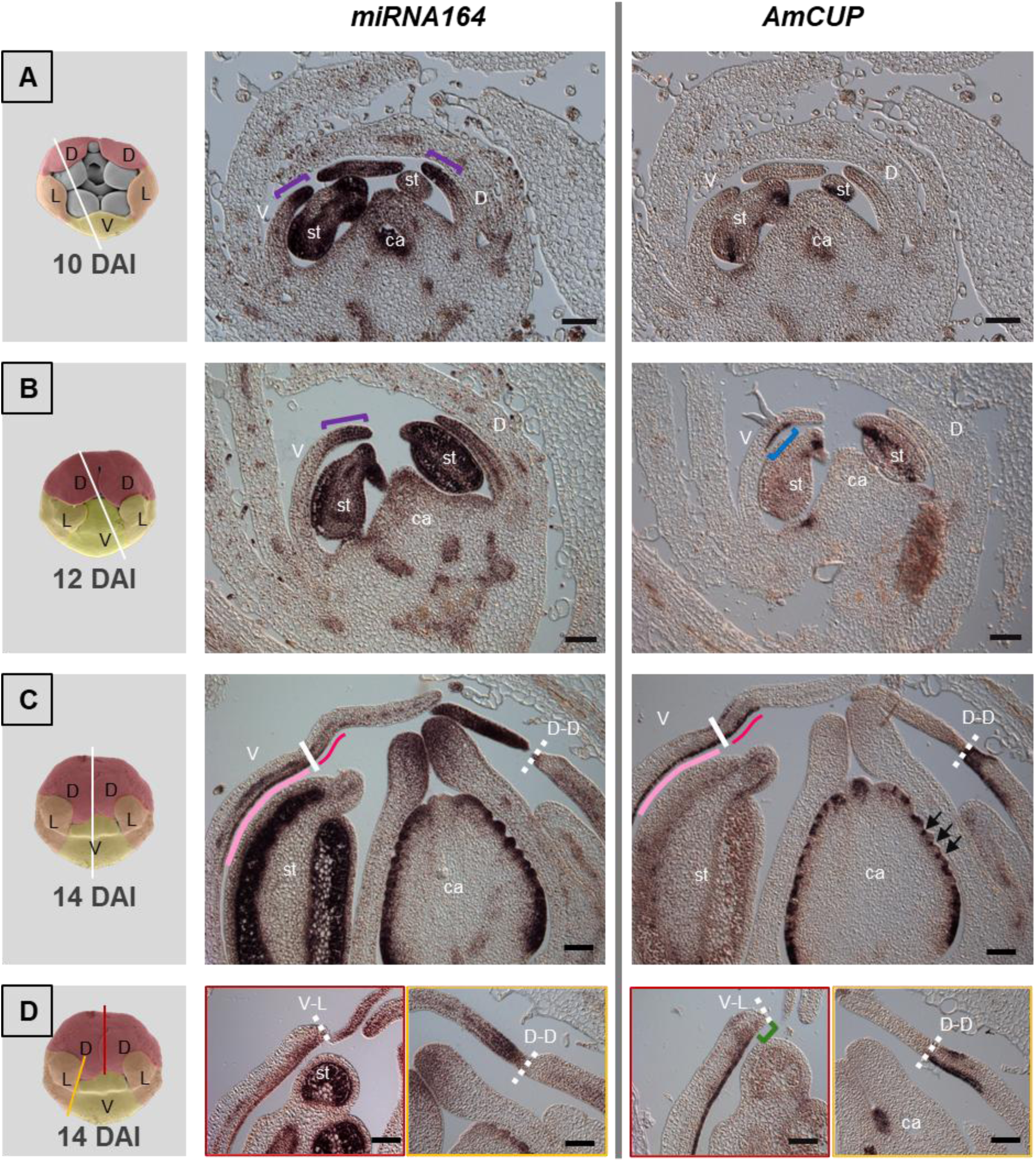
Expression pattern of *miR164* (left) and *AmCUP* (right) at different DAI detected by *in situ* hybridisation. **(A and B)** Expression of *miRNA*, at 10 and 12 DAI was visible in the distal part of the petals (purple brackets) and *AmCUP* was restricted to a small crescent portion of the ventral petal (blue bracket). **(C)** Later, at 14 DAI, both genes co-located in the palate and lip region (light pink and dark pink, respectively), separated by the rim (white line). *miRNA164* was expressed in the stamens and carpel while *AmCUP* seemed to be restricted to the ovary boundaries (black arrows). **(D)** The *miR164* was expressed in the V-L junction (dotted line) while *AmCUP* expression was absent (green bracket). In the dorsal petals *AmCUP* was restricted to the proximal region adjacent to the D-D junction (dotted line) while *miR164* reached the distal region. D (dorsal); V (ventral); st (stamen); ca (carpel). Scale 100 µm.

For *AmBZR1*, at 7 DAI, the expression of *AmBZR1* was observed in the organ primordia, whereas *AmCUP* was expressed at organ boundaries (Fig. 9a). Similar to *miR164*, the expression of *AmBZR1* was first observed at the distal part of the petals (Fig. 9b purple bracket) and then in the V-L junction (Fig. 9c, green bracket). *AmCUP* was expressed in the distinct crescent shape within the lower petal, partially overlapping with *AmBZR1* (Fig. 9c, blue bracket). This overlapping subsequently extended to the palate and lip during a later stage of development (Fig. 9d). Later, *AmBZR1* was expressed distally at the D-D junction, whereas *AmCUP* was expressed proximally (Fig. 9d). *AmBZR1* expression was observed in the distal region of the carpel and in ovules, and *AmCUP* was again limited to the ovule boundaries (Fig. 9d, black arrows).

**Figure 9.**
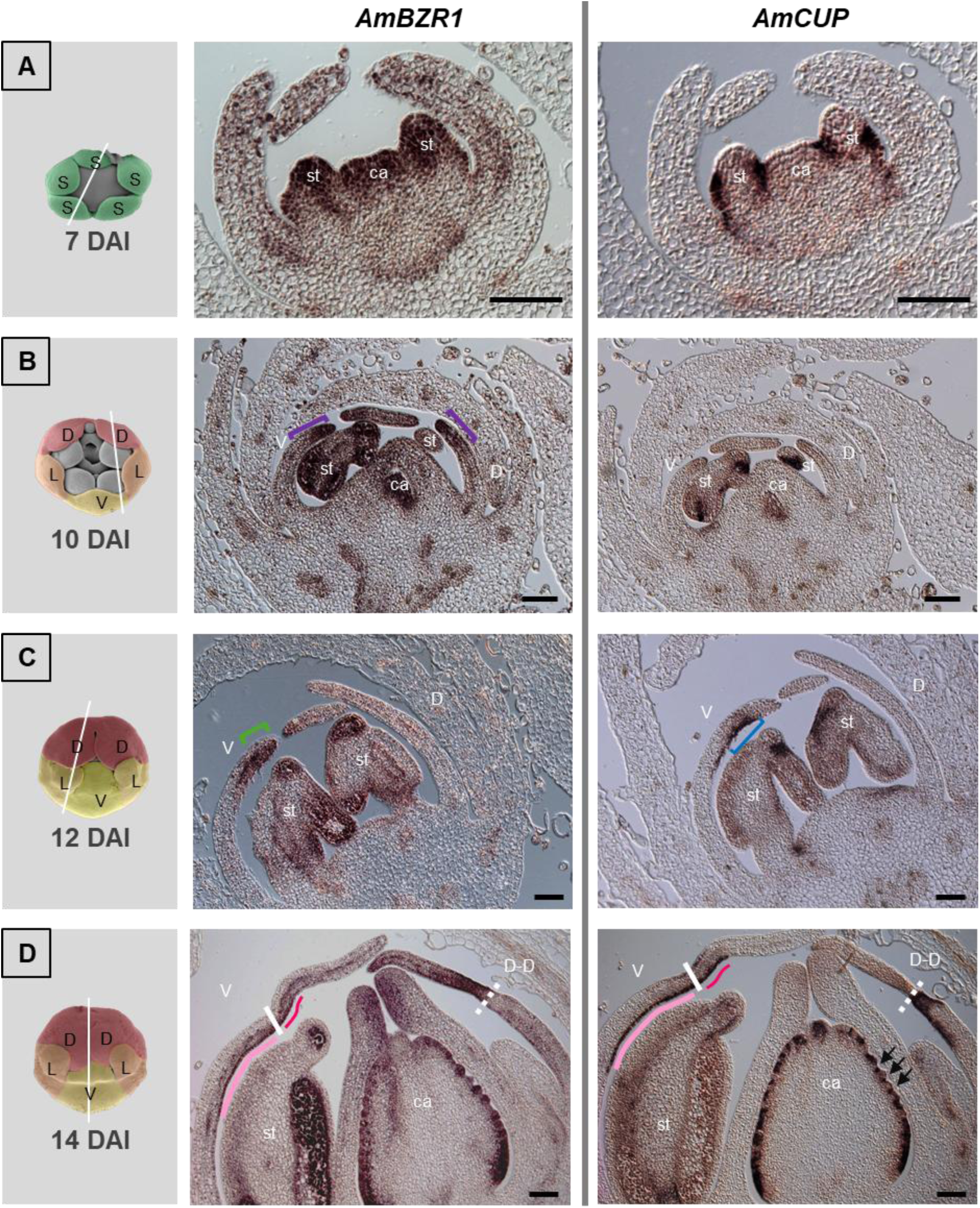
Expression pattern of *AmBZR1* (left) and *AmCUP* (right) at different DAI detected by *in situ* hybridisation. **(A)** Expression of *AmBZR1* at 7 DAI was visible in the centre of organ primordia while *AmCUP* was restricted to the boundaries. **(B)** At 10 DAI *AmBZR1* was expressed at the distal part of petals (purple brackets). **(C)** *AmCUP* was restricted to a small crescent portion of the ventral petal (blue bracket), while *AmBZR1* was expressed in the V-L junction (green bracket). **(D)** At 14 DAI both genes co-localised in the palate and lip region (light pink and dark pink, respectively), separated by the rim (white line). In the dorsal petals *AmCUP* was restricted to the proximal region adjacent to D-D junction (dotted line) while *AmBZR1* was restricted to the distal region. *AmBZR1* is expressed in the stamens and carpel whereas *AmCUP* was expressed in the ovary boundaries (black arrows). D (dorsal); V (ventral); st (stamen); ca (carpel). Scale 100 µm.

*AmNGAL1* was identified as a more dorsally expressed gene in the WT petal samples (Fig. 3). However, its potential role in controlling *AmCUP* led to further analysis through ISH. By 10 DAI, *AmNGAL1* was expressed at organ boundaries, in conjunction with *AmCUP* (Fig. 10a, white arrows). At 12 DAI their expression continued to be co-localised in the palate and ovule boundaries, but not in the lip where *AmNGAL1* was absent (Fig. 10b). Both genes were absent in the V-L junction (Fig. 10c, green bracket) and, while *AmCUP* was expressed at the D-D junction, *AmNGAL1* was expressed at the most lateral part of the dorsal petals (Fig. 10d).

**Figure 10.**
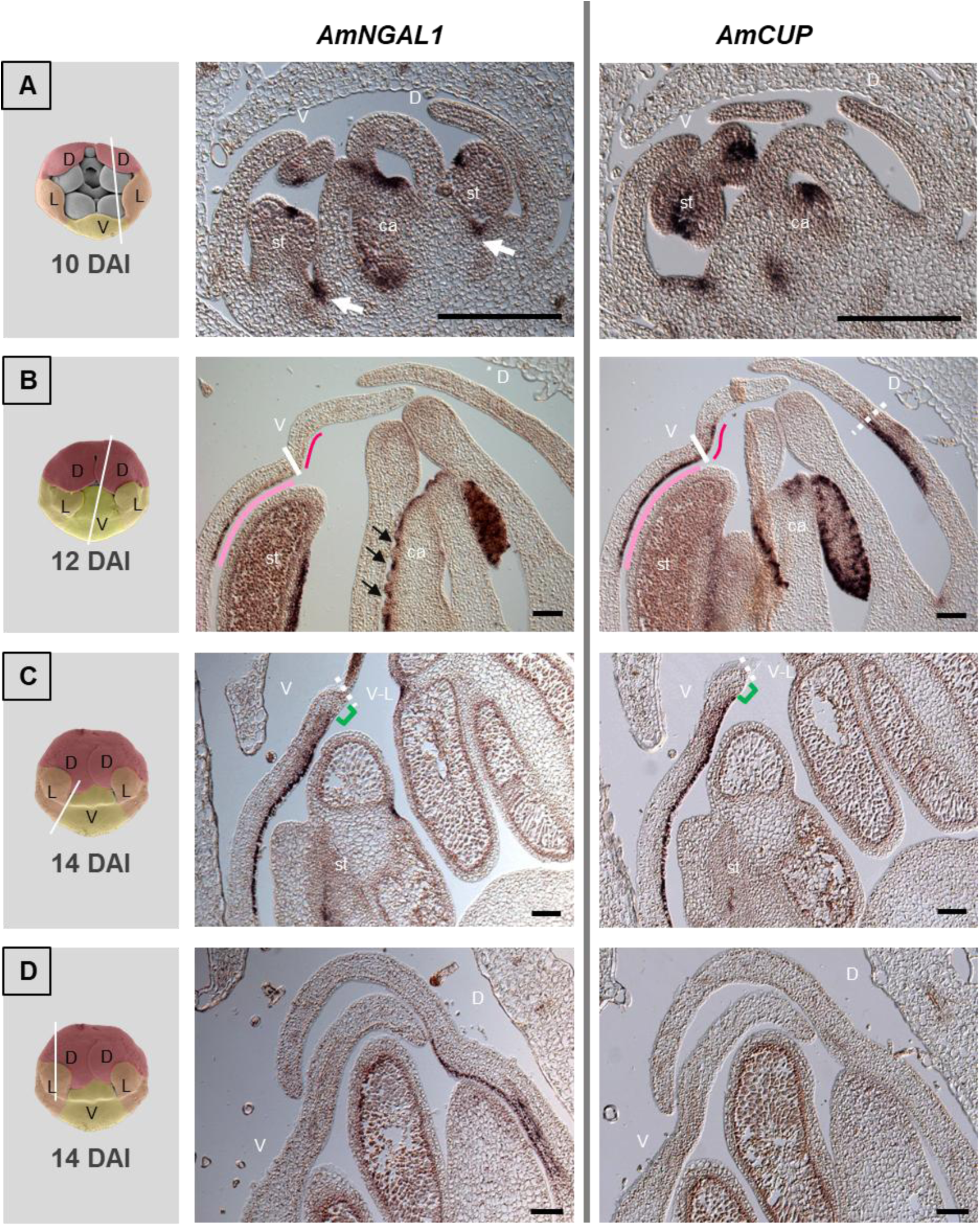
Expression pattern of *AmNGAL1* (left) and *AmCUP* (right) at different DAI detected by *in situ* hybridisation. **(A)** Expression of *AmNGAL1* at 10 DAI overlapped with that of *AmCUP* at organ boundaries (white arrows). **(B and C)** Expression of both genes also overlapped at 12 and 14 DAI, in the palate but not in the lip (light pink and dark pink, respectively, separated by the rim, white line). In the V-L junction (dotted line) *AmCUP* and *AmNGAL1* expression was absent (green bracket). **(D)** *AmNGAL1* was expressed at a more lateral part of the dorsal petals. D (dorsal); V (ventral); st (stamen); ca (carpel). Scale 100 µm.

## Discussion

In *Antirrhinum* flowers, dorsoventral asymmetry arises from the asymmetric action of *AmCYC*, *AmDICH*, *AmRAD* and *AmDIV* along the dorsoventral axis (Luo et al., 1996; Almeida et al., 1997; Luo et al., 1999; Galego and Almeida, 2002; Corley et al., 2005; Costa et al., 2005). Additionally, *AmDIV* and *AmCUP* have been identified as crucial players in the development of the complex-shaped corolla, which depends on differential tissue growth along the proximal-distal axis (Galego and Almeida, 2002; Rebocho et al., 2017a). In the current study, we identified and characterized additional genes putatively involved in the regulatory network controlling flower shape development in *Antirrhinum* (Fig. 11).

**Figure 11.**
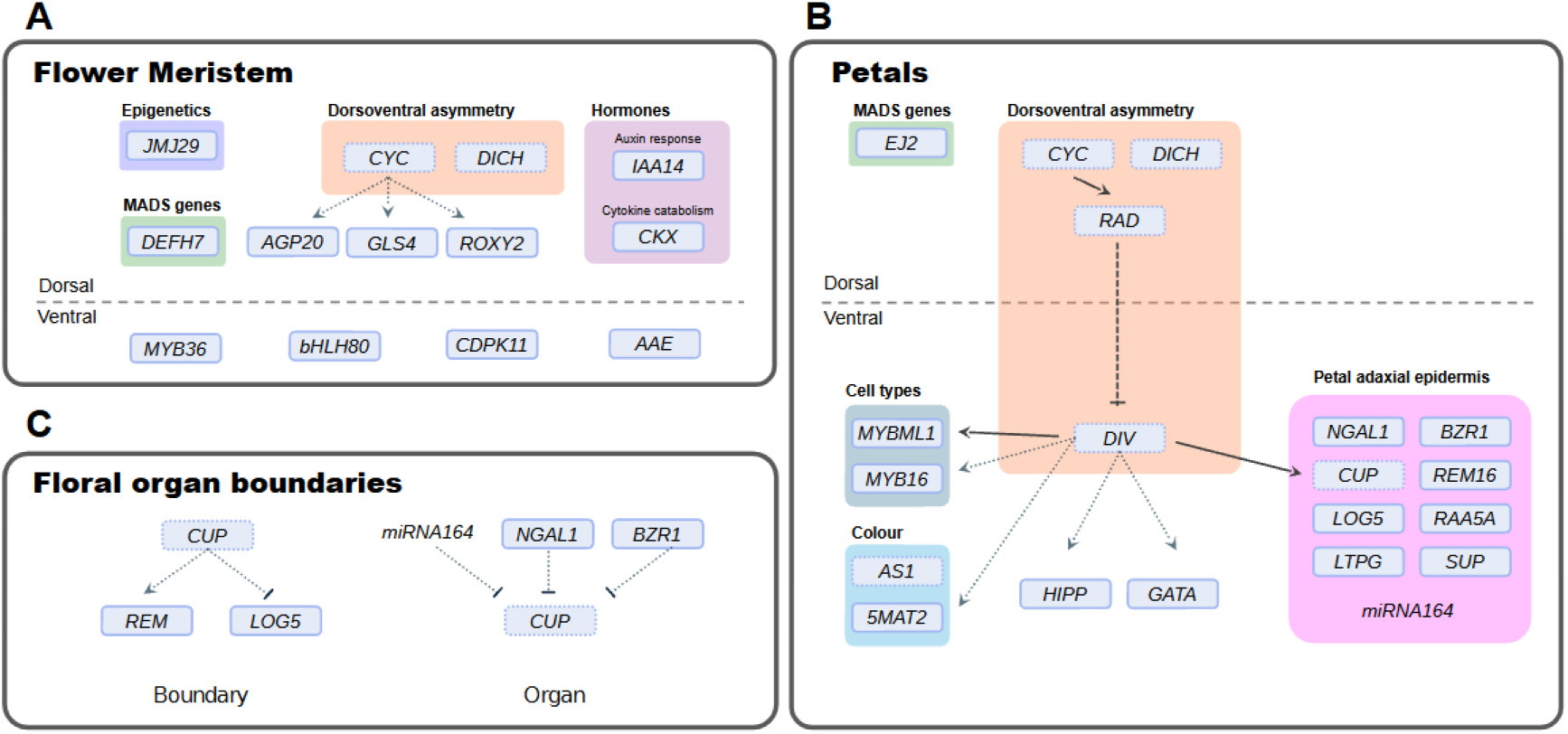
Gene regulatory network controlling *Antirrhinum* flower development. **(A)** In the dorsal regions of the flower meristem, *AmCYC* and *AmDICH* are known to be dorsally expressed establishing dorsoventral asymmetry. Using transcriptomic analysis new genes were identified as more dorsally expressed including members of the *MADS-box* family and genes related to hormones and epigenetic modifications. New putative direct targets of AmCYC (dashed arrows) were identified, namely homologues of *AGP20*, *ROXY2* and *GLS4*. **(B)** During petal development more genes were identified as ventrally expressed. Some of the more ventrally expressed genes were restricted to the adaxial epidermis likely influencing the final shape of the petal. *AmDIV* appears to be a pivotal regulator of genes involved in diverse processes, including petal shape, the production of distinct cell types and pigmentation. **(C)** The expression of *AmCUP* is restricted to organ boundaries by a module that may include *miRNA164*, *AmNGAL1* and *AmBZR1*. At organ boundaries, *AmCUP* may repress the growth of these regions by regulating *AmREM16* and *LOG* genes. Full connection lines represent transcriptional regulations previously described in the literature, whereas dotted connection lines are proposed transcriptional regulation described in this work.

Transcriptomic analysis of two stages of flower development: dorsal and ventral regions of the FM (5 DAI) and dorsal and ventral petals (5 mm wide buds) revealed major differences in DEGs, with *AmCYC* and *AmDICH* being the only genes that showed significant differential expression at both stages. The AmCYC target, *AmRAD* was only upregulated in the dorsal petal of the later stage of flower development analysed. Previous research has shown that the expression of *AmCYC* and *AmDICH* is evident from the initial phase of flower meristem development (Luo et al., 1996; Luo et al., 1999), whereas the dorsal expression of *AmRAD* is observed at a slight later stage (Corley et al., 2005; Costa et al., 2005). In the dorsal petals, AmRAD limits AmDIV protein activity to regulate dorsal petal identity (Galego and Almeida, 2002; Raimundo et al., 2013). Surprisingly, another gene from the *RAD* subfamily, *RADIALIS-like 6* (*RADL6*), was found to be more expressed in the ventral petals. It is possible that the *RADL6* homologue serves other roles and may not participate in the antagonistic module that regulates *AmDIV* action.

A greater number of genes were found to be more expressed in the dorsal region of the FM than in the ventral, whereas the reverse was observed in the petal samples. This suggests that, at an early stage, more genes with asymmetric expression are required to establish and/or maintain the developmental programme of the dorsal domain of the FM. Later, during the development of the more complex-shaped ventral petals, which contain a greater number of cell types, (Glover et al., 1998; Perez-Rodriguez et al., 2005), have a yellow pigmentation and form a characteristic wedge shape, a more complex genetic network may be required. This temporal difference, with a higher number of more dorsally expressed genes early in development and a higher number of more ventrally expressed genes later, is also found in *S. speciosa* and *P. sativum* (Jiao et al., 2017; Pan et al., 2022).

### Dorsally expressed genes related to floral meristem organisation

At an early stage, *AmCYC* plays an essential role in flower development by inhibiting the growth of dorsal regions of the FM and influencing petal number (Luo et al., 1996). Our transcriptomic analysis of early meristem development identified 29 genes more dorsally expressed. Among these were homologues of genes from *Arabidopsis* involved with floral meristem organisation *ROXY2* and cytokinin dehydrogenase *CKX* (Fig. 11a). In *Arabidopsis*, *roxy1* mutants exhibit a reduction in the number of petals and display abnormal petal bending, phenotypes that can be rescued by its homologue *ROXY2* (Xing et al., 2005; Xing and Zachgo, 2008). By contrast, *Arabidopsis ckx3 ckx5* double mutants exhibit an increase in the number of floral organs and have a larger petal surface area (Bartrina et al., 2011). In *Antirrhinum*, both *ROXY2* and *CKK* homologues could be involved in the establishment of the number of floral organs and petal development with *ROXY2* potentially acting downstream of AmCYC, as it contains AmCYC binding sites in its promoter. A callose synthase, a *GSL4* homologue, was identified as more dorsally expressed and also contained AmCYC binding sites in its promoter. Mutants in *Arabidopsis* with impaired callose synthesis are affected in cell division, leading to defects in tissue organisation, cell number, and division plane (Chen et al., 2009; Thiele et al., 2009). Callose deposition in plasmodesmata can also create asymmetric auxin gradients necessary for *Arabidopsis* hypocotyl growth response (Han et al., 2014). Asymmetric auxin gradients in the FM may influence *AmCYC* expression, given that auxin has been suggested to influence petal identity (Bergbusch, 1999). *AmCYC* may also be involved in cell division and/or creating auxin gradients in the dorsal petal through the regulation of callose synthesis genes.

Another more dorsally expressed gene in the FM, *AmDEFH7,* belongs to an ancient *TM8* lineage of MIKC^C^-type MADS-box genes (Li, 2002; Coenen et al., 2018). In *Nicotiana benthamiana*, the *NbTM8* homologue plays a role in both phase transition and the determination of floral organ number in the outer three whorls (Coenen et al., 2018). In *Antirrhinum*, the tissues showing the greatest *AmDEFH7* expression are the stem and flower, yet *AmDEFH7* function remains elusive (Coenen et al., 2018). Regulators of early *AmCYC* expression are still unknown, yet late expression is maintained by the B-class MADS-box gene *AmDEF* (Clark and Coen, 2002). One hypothesis that requires further study is that *AmDEFH7* may also regulate *AmCYC* expression during the initial stages of FM development.

### Dorsoventral genes are also modulated along the proximal-distal axis

Transcriptome analysis of the *Antirrhinum* petals showed a greater difference in gene expression between dorsal and ventral petals than between the dorsal and ventral regions of the FM (Fig. 11b). The comparison between the two petal types identified 37 genes more expressed in dorsal petals and 68 more expressed in the ventral petals. During the formation of the ventral petal, a furrow is formed which later expands giving rise to the palate and lip regions, that bend at the rim (Green et al., 2010; Rebocho et al., 2017a; Rebocho et al., 2017b). Dorsoventral gene activity, together with local factors, modulates tissue growth rates and pattern, consequently shaping the corolla. This is exemplified by *AmDIV* action, which promotes the expression of *AmCUP* in the palate-lip region and leads to differential tissue growth of regions of the ventral petal (Rebocho et al., 2017a). A hypothesis is that genes identified as more ventrally expressed could also be regulated along the petal proximal-distal axis and thus influence the shape of the ventral petal. To test this hypothesis, the expression of the 68 genes previously identified as more ventrally expressed was analysed across the four regions of the ventral petal, namely the tube, palate, lip and lobe. This analysis generated five clusters that included genes expressed in: lobe-lip, palate, palate-tube, lip-palate and tube. Among genes more expressed in the lip-palate were *AmCUP* and the homologues of *LBD36* and *GSO1.* In *Arabidopsis*, *LBD36* together with its closest homologue *ASYMMETRIC LEAVES2 (AS2)*, control proximal-distal polarity and epidermal cell fate determination during petal development (Chalfun-Junior et al., 2005). *GSO1* is involved in the longitudinal expansion of epidermal cells in the hypocotyl of *Arabidopsis* (Tsuwamoto et al., 2008). The expression of a *LBD36* and *GSO1* homologues may be necessary to maintain differential growth between abaxial and adaxial sides of the petal to achieve the curvature associated with the palate-lip region. The adaxial epidermis of the palate-lip region has increased cell proliferation and expression of cell cycle markers contributing to the petal curvature (Gaudin et al., 2000; Crawford et al., 2004).

The expression of genes from other clusters was then analysed by ISH including *LOG5* from the lobe-lip cluster, *RAA5a* from the palate and *SUP* and *LGPT13* from the palate-tube. These genes were expressed mainly in the adaxial epidermis in the middle of the ventral petal, indicating a differential regulation of adaxial-abaxial expression along the proximal-distal axis of the ventral petal. The transcriptomic analysis detected the presence of *LOG5* expression in the lobe-lip regions; however, ISH analysis revealed that *LOG5* expression was not limited to these regions, being also observed in the palate. This is possibly due to the different developmental times of sampling between the two analyses, suggesting that in addition to its ventral expression, *LOG5* may also be subject to regulation across the proximal-distal axis according to developmental stages. *LOG* genes play a crucial role in the direct synthesis of biologically active cytokinins, which promote cell division (Riou-Khamlichi et al., 1999; Kurakawa et al., 2007; Kuroha et al., 2009; Tokunaga et al., 2012; Yang et al., 2021). A *LOG* gene has been recently identified in *Antirrhinum*, as more expressed in the ventral tube than its dorsal counterpart, possibly being involved in the development of the curve-shaped gibba tube region (Cullen et al., 2023). The *LOG5* identified here, may have been recruited to shape the curved ventral furrow by modulating cytokinin levels and promoting cell division in the adaxial epidermis.

*LTPGs* are involved in cuticular wax export and are present in rapidly growing epidermal cells, capable of responding to changes in cell geometry (DeBono et al., 2009; Ambrose et al., 2013). The Rab family of GTPases regulates membrane trafficking, and *RAB*-*A5c* is essential for maintaining correct anisotropic cell growth during organogenesis in *Arabidopsis* (Kirchhelle et al., 2016). In *Antirrhinum*, *LTPG13* and *RAA5a* homologues may play roles in the development of the palate-tube and palate regions, respectively, and in the maintenance of the correct growth of epidermal cells specific to these areas. The boundary gene *SUP* from *Arabidopsis* controls FM size and floral organogenesis (Bowman et al., 1992; Sakai et al., 1995; Xu et al., 2018). At the whorl 3/4 boundary (stamen/carpel), *SUP* represses the auxin biosynthesis genes *YUCCA1/4*, which terminates stem cell activity (Xu et al., 2018). In *Medicago truncatula*, *MtSUP* regulates the number of organs in the inner whorls 3/4, as well as in the petal whorl (Rodas et al., 2021). It is possible that in *Antirrhinum*, as in *Medicago*, a *SUP* homologue extends its action to the petal whorl maintaining boundaries between different regions of the ventral petal. However, this action requires further elucidation as its expression in the ventral petal overlaps with that of auxin biosynthesis gene *AmYUCCA1 (*Rebocho et al. 2017a).

The ventral petal of *Antirrhinum* is also distinguished by its colour and regions of high density of trichomes. Genes associated with anthocyanin biosynthesis exhibited higher expression in lip and lobe regions, characterised by a more intense magenta colour, whereas the aurone biosynthesis genes were more expressed in the tube and palate, where the yellow pigmentation is visible. The expression of *5MAT2* and *5MAT1* homologues, which catalyse anthocyanin malonylation (Suzuki et al., 2001; Suzuki et al., 2004), in the tube and palate-lip, respectively, raises the hypothesis of specific pigment production in these regions. In *Antirrhinum* the *MYBML* family, comprising *AmMIXTA* and *AmMYBML1-3*, is involved in cell specification in the petal epidermis, with only *AmMYBML1* initiating trichomes (Perez-Rodriguez et al., 2005). Here we showed that genes of the *MYBML* family (*AmMYBL1*, and homologues of *MYB160* and *MYB16*) also seem to be regulated along the proximal-distal axis in ventral petals and we also identified a more ventrally expressed new member of the *MYBML* family, closely related to *AmMYBML1*, which is likely to be the result of a recent duplication. This suggests that more genes may be involved in cell specification and trichome initiation and/or branching in *Antirrhinum*.

### *AmDIV* influences different aspects of the ventral petal development

*AmDIV* plays a central role in the development of various characteristics of the ventral petal, but so far only *AmCUP* was identified as a direct target (Rebocho et al., 2017a). Here, 26 new putative *AmDIV* targets were identified by selecting genes that were more expressed in the ventral petal and ventralised *cyc dich* mutant, and simultaneously downregulated in *cyc dich div* mutants. These genes included homologues of *SUP, RAA5a,* and *REM16*, whose expression in the ventral adaxial epidermis, overlaps with *AmDIV* expression. *AmMYBML1* was also identified as potentially regulated by *AmDIV.* This agrees with the findings of Perez-Rodriguez et al. (2005), that demonstrated a reduction in *AmMYBML1* expression in *div* mutants.

The promoters of homologues of *GATA9*, *HIPP22*, *MYB16* and *5MAT2*, contained DIV-binding sites within the first 1 kb, suggesting a possible direct regulation by AmDIV (Fig. 11b). The expression of *GATA9* was higher in the palate-lip region. In *Arabidopsis*, the *HANABA TARANU* gene, a member of the *GATA* family, has been implicated in establishing organ boundaries by modulating cytokinin homeostasis, and regulating floral organ development (Ding et al., 2015; Leonte, 2020). Genes from the *HIPP* family, act as negative regulators of flowering time in *Arabidopsis* (Zschiesche et al., 2015), and the lack of *HIPP32-34* genes results in defects in leaf and petal organogenesis, leading to smaller and narrower petals (Leonte, 2020). The presence of *DIV*-binding sites in the *5MAT2* and *MYB16* promoters suggests that AmDIV may have a direct involvement in pigment production and trichome development processes. Genes involved in trichome development and anthocyanin biosynthesis are often regulated by complexes between MYB, bHLH and WD40 proteins (Wang et al. (2021b), Yan et al. (2021) and Li et al. (2022). Overall, *AmDIV* appears to coordinate a complex network of genes that influence petal shape, anthocyanin pigmentation and trichome development during ventral petal development.

### *AmCUP* role in hormonal regulation is likely region specific

*AmCUP* is involved in repressing growth at organ boundaries and later promoting growth in the ventral furrow (Rebocho et al., 2017a). *AmCUP* possibly modulates auxin levels through the auxin biosynthesis gene *AmYUCCA1,* which may enhance growth in the ventral petal, but not in the organ boundaries (Rebocho et al., 2017a). A transcriptomic analysis was done with the *cup* mutant flowers to better understand the role of *AmCUP* in flower development. A larger number of genes was identified as up regulated in the mutant suggesting a possible repressive role for *AmCUP*. This is supported by the fact that the AmCUP protein contains a NARD subdomain responsible for repressing transcription (Hao et al., 2010). Transcriptome analysis revealed several DEGs related to auxin. Two of these genes were homologues of the auxin transporters *TRN2* and *PIL*, up and down regulated in the mutant, respectively. In *Arabidopsis*, *TRN2* is involved in the generation of auxin maxima that affects gynoecium formation and leaf shape (Cnops et al., 2006; Yamaguchi et al., 2017; Yamaguchi et al., 2018). *PILs* function as intracellular transporters of auxin to the endoplasmic reticulum, limiting the rate of available auxin in the nucleus (Barbez et al., 2012). The *cup* mutant showed an ambiguous pattern of auxin signalling, with a *ARF* homologue downregulated and two *AUX/IAAs* being upregulated. Responses to changes in auxin levels are mediated by auxin early response genes like Aux/IAA and ARFs (Luo et al., 2018; Cancé et al., 2022). Some of these proteins function in a module where in low auxin conditions, Aux/IAA represses ARFs, whereas in the presence of auxin, Aux/IAA are degraded, releasing ARFs repression and activating transcription (Salehin et al., 2015; Israeli et al., 2020; Cancé et al., 2022). The role of *AmCUP* in the regulation of auxin-related genes requires further investigation, as it may be spatially dependent. Nevertheless, it seems plausible that auxin biosynthesis is likely impaired in the *cup* mutant, as evidenced by the downregulation of *YUCCA6* homologue in the mutant, in addition to *AmYUCCA1* reported by Rebocho et al. (2017a).

The transcriptome analysis also showed an up-regulation of *ACS*-*like*, *GA2OX1* and *GA2OX8* genes, suggesting an involvement of *AmCUP* in the inhibition of ethylene biosynthesis and gibberellin catabolism. In gerbera and rose, ethylene regulates petal size by inhibiting cell expansion (Ma et al., 2008; Huang et al., 2020; Cheng et al., 2021). In *Rosa hybrida,* ethylene control of cell expansion seems to involve a module consisting of the *miRNA164* and the NAC gene *RhNAC100* Pei et al., 2013). Gerbera petal size is also controlled by gibberellins, specifically a *GASA-like* gene that promotes elongation (Li et al., 2015b; Jiang et al., 2022). *AmCUP* inhibition of growth at organ boundaries may be mediated in part by a reduction in ethylene and gibberellins, which inhibit cell expansion. The down regulation of *LOG1* and *LOG3* homologues genes in *cup* mutants also points for a role of *AmCUP* in the regulation of cytokinin biosynthesis. The expression of *LOG5,* which was studied by ISH, overlaps with *AmCUP* in the ventral petal. However, in earlier stages of development *LOG5* is expressed in the centre of the organ primordia (Supplementary Figure 5). A potential mechanism for *AmCUP* growth repression at organ boundaries may involve the modulation of cytokinin levels through the action of *LOG* genes. The apparent antagonistic effects of *AmCUP* on hormonal regulation suggests a region-specific role. *AmCUP* may inhibit growth in organ boundaries by reducing levels of auxin, ethylene, gibberellin and cytokinin, whereas in the ventral furrow, due to the presence of different factors, *AmCUP* may promote growth by elevating the levels of these hormones.

The gene *AmREM16*, downregulated in the *cup* mutant, was identified as possibly being regulated by *AmDIV*. ISH revealed that *AmREM16* expression in petal and ovule boundaries is similar to that of *AmCUP,* and in the ventral petal *AmREM16* expression overlaps with that of both *AmCUP* and *AmDIV*. *AmREM16* belongs to the B3 domain *REPRODUCTIVE MERISTEM* family of transcription factors, with its closest homologue in *Arabidopsis* being *REM16* (Swaminathan et al., 2008; Romanel et al., 2009). In *Arabidopsis*, some genes of this family have been associated with the development of the female gametophyte (Matias-Hernandez et al., 2010; Mendes et al., 2016; Caselli et al., 2019), initiation of ovule primordia (Gomez et al., 2018), flowering time (Levy et al., 2002; Heo et al., 2012; Richter et al., 2019; Yu et al., 2020) and transition to the reproductive phase (Manrique et al., 2023). In *Antirrhinum*, *AmREM16* potentially controls organ boundaries and ventral petal shape, likely acting downstream of *AmDIV* and *AmCUP*.

### *AmCUP* is possibly regulated by *miR164*, *AmBZR1* and *AmNGAL1*

In the ventral furrow *AmDIV* promotes *AmCUP* expression, yet *AmCUP* expression in petal junctions seems to be independent of *AmDIV* (Rebocho et al., 2017a). This hinted to the existence of more *AmCUP* regulators specially at organ boundaries. When analysing the list of DEGs between dorsal and ventral petals, two genes were identified as possible *AmCUP* regulators, one more ventrally expressed, *AmBZR1*, and another more dorsally expressed, *AmNGAL1*. In *Arabidopsis*, these two genes regulate the homologues of *AmCUP,* known as *CUCs* (Gendron et al., 2012a; Shao et al., 2020). In various species, the *miR164* is also known for its pivotal role in regulating *AmCUP* homologues and defining organ boundaries (Laufs et al., 2004; Mallory et al., 2004; Baker et al., 2005; Berger et al., 2009; Hendelman et al., 2013; Gonçalves et al., 2015; Zheng et al., 2019; Wang et al., 2021a). To test the hypothesis of a conserved mechanism of *AmCUP* regulation, an ISH for *AmBZR1*, *AmNGAL1* and *miR164* was conducted. In the petals, the expression patterns of *miR164* and *AmCUP* are generally complementary, except in the ventral furrow where they overlap. Complementarity in expression at organ boundaries, dorsal petals and early stages of the ventral petal suggests a potential role for *miR164* to repress *AmCUP* in these regions. This repressive action is further supported by the presence of conserved *miR164* regulatory sites in *AmCUP* (Hao et al., 2010; Vialette-Guiraud et al., 2016). The later overlap in expression of *AmCUP* and *miR164* in the ventral petal could be influenced by the presence of other genes. The presence of *AmDIV* promotes *AmCUP* expression and may counteract the repressive effect of *miR164*. The regulatory mechanism of the *miRNA/CUP* module differs among species. In *Arabidopsis*, a *miR164* regulates the spatial distribution of *CUCs* and their transcript accumulation (Baker et al., 2005; Nikovics et al., 2006; Raman et al., 2008), whereas in tomato, *miR164* and *SlNAM* expression are complementary, indicating an exclusivity mechanism (Berger et al., 2009). The *miR164/AmCUP* regulation in *Antirrhinum* potentially employs a hybrid system where expression is exclusive at the boundary locations, but in the palate-lip region, the presence of other genes may not affect or only lead to a reduction in the expression of *AmCUP* (Fig. 11c). Expression of *miR164* was also detected at the tips of the carpels, where *AmCUP* is absent. In *Arabidopsis*, carpel fusion is also controlled by *CUC1-CUC2/miR164*, as well as the formation of leaf serration patterns (Nikovics et al., 2006; Vialette-Guiraud et al., 2011; Kamiuchi et al., 2014; Vialette-Guiraud et al., 2016).

The expression of *AmBZR1* was similar to that of *miR164*, with *AmBZR1* also exhibiting a complementarity with *AmCUP* expression, except in the palate-lip region. *BZR1* has a direct repressive effect on *CUC* genes at boundaries in the *Arabidopsis* shoot apical meristem (Gendron et al., 2012a). Brassinosteroid signalling can also activate a *BZR1* homologue, *BRI1-EMS-SUPPRESSOR 1* that, together with *TOPLESS*, repress *CUC3* in boundary regions (Espinosa-Ruiz et al., 2017). The complementary expression of *AmBZR1* and *AmCUP* in the FM, suggests that, as with *Arabidopsis*, *AmBZR1* may serve to limit *AmCUP* to the organ boundaries, and later to ovule boundaries (Fig. 11c). *BZR1* and *CUC2* are positive regulators of ovule initiation, yet they act in an independent manner (Gendron et al., 2012a; Barro-Trastoy et al., 2022). In *Arabidopsis*, *BZR1* directly binds to the BR-response element (CGTG(T/C)G) sequence (He et al., 2005; Gampala et al., 2007). In the *AmCUP* promoter, the binding site for *BZR1* is present 4 times within the 1 kb before the ATG codon (Supplementary Table 4), reinforcing the hypothesis of a potential direct regulation.

*AmNGAL1*, was identified by transcriptomic analysis as more expressed in dorsal petals, a pattern similarly observed in *S. speciosa* (Pan et al., 2022). ISH analysis confirmed the expression in dorsal petals but also revealed *AmNGAL1* expression at organ boundaries and ventral petals. Large expression domains can mask smaller, yet possibly important expression patterns. *AmNGAL1* expression largely overlapped with that of *AmCUP* except in the dorsal petal. A similar expression overlap is observed in *Arabidopsis*, between *NGAL3* and *CUC2,* where *NGAL3* represses *CUC2* in leaf margins to form serration patterns (Engelhorn et al., 2012). Moreover, *NGAL1* is also involved in the formation leaf serration patterns by directly repressing *CUC2* (Shao et al., 2020). *NGAL2/3* repress *CUC* genes during the establishment of boundaries in cauline axillary meristems (Hibara et al., 2006; Nicolas et al., 2022). The loss of *NGALs* enhances the phenotypes of *cuc* mutants, including the development of an unfused gynoecium and the presence of enhanced leaf serrations (Engelhorn et al., 2012; Shao et al., 2020). It is possible that regulatory modules involved in *Arabidopsis* leaf morphology, including *CUCs, NGALs* and *miRNA164*, might be conserved in *Antirrhinum* as part of the regulatory network controlling floral organ boundaries. The *CINCINNATA* in *Antirrhinum*, has been shown to repress growth in leaf margins and promote growth in petals (Nath et al., 2003; Crawford et al., 2004), demonstrating a direct link between the development of both leaves and petals. *AmNGAL1*, *AmBZR1* and *miR164* may be involved in the establishment of organ boundaries by repressing *AmCUP.* The precise function of these factors in the formation of the ventral petal remains uncertain due to their expression overlapping with that of *AmCUP*.

In conclusion, this study has utilised transcriptomic data supported by ISH to deepen our understanding of how asymmetric gene expression along the dorsoventral and proximal-distal axis may shape complex flowers of *Antirrhinum*. During early FM development, genes that are more expressed in the dorsal region include those involved in hormonal regulation. Subsequently, genes involved in various aspects of the ventral petal development, including shape, pigmentation and the differentiation of specific cell types, are likely to be subject to *AmDIV* regulation. Genes identified as more ventrally expressed are also differentially expressed along the proximal-distal axis, likely regulating the development of different regions of the ventral petal. These results extend the idea that genes involved in dorsoventral asymmetry interact with genes along the proximal-distal axis to contribute to the formation of a complex corolla shape (Green et al., 2010; Rebocho et al., 2017a; Rebocho et al., 2017b). Genes with asymmetric expression along the dorsoventral axis have also been implicated in the establishment of floral organ boundaries in relation to *AmCUP*. In particular, the regulation of *AmCUP* at the floral boundaries may involve similar mechanisms to those regulating primordia outgrowth in *Arabidopsis*. Overall, the data provide a valuable resource for further understanding the intricate genetic interactions that influence the development of complex dorsoventral asymmetric flowers.

## Acknowledgements

We would also like to acknowledge the contribution of Mike Scanlon, whose expertise in the laser microdissection technique has significantly advanced our research.

## References

1. Aida M, Ishida T, Fukaki H, Fujisawa H, Tasaka M (1997) Genes involved in organ separation in Arabidopsis: an analysis of the cup-shaped cotyledon mutant. Plant Cell 9: 841–857

2. Albert NW, Butelli E, Moss SMA, Piazza P, Waite CN, Schwinn KE, Davies KM, Martin C (2021) Discrete bHLH transcription factors play functionally overlapping roles in pigmentation patterning in flowers of Antirrhinum majus. New Phytologist 231: 849– 863

3. Almeida J, Rocheta M, Galego L (1997) Genetic control of flower shape in Antirrhinum majus. Development 124: 1387–1392

4. Altschul SF, Gish W, Miller W, Myers EW, Lipman DJ (1990) Basic local alignment search tool. J Mol Biol 215: 403–410

5. Ambrose C, DeBono A, Wasteneys G (2013) Cell Geometry Guides the Dynamic Targeting of Apoplastic GPI-Linked Lipid Transfer Protein to Cell Wall Elements and Cell Borders in Arabidopsis thaliana. PLoS One 8: e81215

6. Anders S, Pyl PT, Huber W (2015) HTSeq-A Python framework to work with high-throughput sequencing data. Bioinformatics 31: 166–169

7. Andrews S (2023) FastQC: A quality control tool for high throughput sequence data. v0.12.1, https://github.com/s-andrews/FastQC

8. Ashburner M, Ball CA, Blake JA, Botstein D, Butler H, Cherry JM, Davis AP, Dolinski K, Dwight SS, Eppig JT, et al (2000) Gene ontology: tool for the unification of biology. The Gene Ontology Consortium. Nat Genet 25: 25–29

9. Bailey TL, Johnson J, Grant CE, Noble WS (2015) The MEME Suite. Nucleic Acids Res 43: W39–49

10. Baker CC, Sieber P, Wellmer F, Meyerowitz EM (2005) The early extra petals1 mutant uncovers a role for microRNA miR164c in regulating petal number in Arabidopsis. Curr Biol 15: 303–315

11. Barbez E, Kubeš M, Rolčík J, Béziat C, Pěnčík A, Wang B, Rosquete MR, Zhu J, Dobrev PI, Lee Y, et al (2012) A novel putative auxin carrier family regulates intracellular auxin homeostasis in plants. Nature 485: 119–122

12. Barro-Trastoy D, Gomez MD, Blanco-Touriñán N, Tornero P, Perez-Amador MA (2022) Gibberellins regulate ovule number through a DELLA–CUC2 complex in Arabidopsis. The Plant Journal 110: 43–57

13. Bartrina I, Otto E, Strnad M, Werner T, Schmülling T (2011) Cytokinin regulates the activity of reproductive meristems, flower organ size, ovule formation, and thus seed yield in Arabidopsis thaliana. Plant Cell 23: 69–80

14. Bateman A, Martin MJ, Orchard S, Magrane M, Ahmad S, Alpi E, Bowler-Barnett EH, Britto R, Bye-A-Jee H, Cukura A, et al (2023) UniProt: the Universal Protein Knowledgebase in 2023. Nucleic Acids Res 51: D523–D531

15. Baumann K, Perez-Rodriguez M, Bradley D, Venail J, Bailey P, Jin H, Koes R, Roberts K, Martin C (2007) Control of cell and petal morphogenesis by R2R3 MYB transcription factors. Development 134: 1691–1701

16. Beld M, Martin C, Huits H, Stuitje AR, Gerats AGM (1989) Flavonoid synthesis in Petunia hybrida: partial characterization of dihydroflavonol-4-reductase genes. Plant Mol Biol 13: 491–502

17. Bergbusch VL (1999) A note on the manipulation of flower symmetry in Antirrhinum majus. Ann Bot 83: 483–488

18. Berger Y, Harpaz-Saad S, Brand A, Melnik H, Sirding N, Alvarez JP, Zinder M, Samach A, Eshed Y, Ori N (2009) The NAC-domain transcription factor GOBLET specifies leaflet boundaries in compound tomato leaves. Development 136: 823–832

19. Bhatia N, Wilson-Sánchez D, Strauss S, Vuolo F, Pieper B, Hu Z, Rambaud-Lavigne L, Tsiantis M (2023) Interspersed expression of CUP-SHAPED COTYLEDON2 and REDUCED COMPLEXITY shapes Cardamine hirsuta complex leaf form. Current Biology 33: 2977–2987

20. Bilsborough GD, Runions A, Barkoulas M, Jenkins HW, Hasson A, Galinha C, Laufs P, Hay A, Prusinkiewicz P, Tsiantis M (2011) Model for the regulation of Arabidopsis thaliana leaf margin development. Proc Natl Acad Sci U S A 108: 3424–3429

21. Blein T, Pulido A, Vialette-Guiraud A, Nikovics K, Morin H, Hay A, Johansen IE, Tsiantis M, Laufs P (2008) A conserved molecular framework for compound leaf development. Science 322: 1835–1839

22. Bowman JL, Sakai H, Jack T, Weigel D, Mayer U, Meyerowitz EM (1992) Superman, a regulator of floral homeotic genes in Arabidopsis. Development 114: 599–615

23. Bradley D, Carpenter R, Sommer H, Hartley N, Coen E (1993) Complementary floral homeotic phenotypes result from opposite orientations of a transposon at the plena locus of Antirrhinum. Cell 72: 85–95

24. Busch A, Zachgo S (2009) Flower symmetry evolution: towards understanding the abominable mystery of angiosperm radiation. BioEssays 31: 1181–1190

25. Cancé C, Martin-Arevalillo R, Boubekeur K, Dumas R (2022) Auxin response factors are keys to the many auxin doors. New Phytologist 235: 402–419

26. Carpenter R, Coen ES (1990) Floral homeotic mutations produced by transposon-mutagenesis in Antirrhinum majus. Genes Dev 4: 1483–1493

27. Caselli F, Beretta VM, Mantegazza O, Petrella R, Leo G, Guazzotti A, Herrera-Ubaldo H, de Folter S, Mendes MA, Kater MM, et al (2019) REM34 and REM35 Control Female and Male Gametophyte Development in Arabidopsis thaliana. Front Plant Sci 10: 1351

28. Chalfun-Junior A, Franken J, Mes JJ, Marsch-Martinez N, Pereira A, Angenent GC (2005) ASYMMETRIC LEAVES2-LIKE1 gene, a member of the AS2/LOB family, controls proximal-distal patterning in Arabidopsis petals. Plant Mol Biol 57: 559–575

29. Chen XY, Liu L, Lee EK, Han X, Rim Y, Chu H, Kim SW, Sack F, Kim JY (2009) The Arabidopsis callose synthase gene GSL8 is required for cytokinesis and cell patterning. Plant Physiol 150: 105–113

30. Cheng C, Yu Q, Wang Y, Wang H, Dong Y, Ji Y, Zhou X, Li Y, Jiang CZ, Gan SS, et al (2021) Ethylene-regulated asymmetric growth of the petal base promotes flower opening in rose (Rosa hybrida). Plant Cell 33: 1229–1251

31. Cheng X, Peng J, Ma J, Tang Y, Chen R, Mysore KS, Wen J (2012) NO APICAL MERISTEM (MtNAM) regulates floral organ identity and lateral organ separation in Medicago truncatula. New Phytol 195: 71–84

32. Clark JI, Coen ES (2002) The cycloidea gene can respond to a common dorsoventral prepattern in Antirrhinum. Plant Journal 30: 639–648

33. Cnops G, Neyt P, Raes J, Petrarulo M, Nelissen H, Malenica N, Luschnig C, Tietz O, Ditengou F, Palme K, et al (2006) The TORNADO1 and TORNADO2 genes function in several patterning processes during early leaf development in Arabidopsis thaliana. Plant Cell 18: 852

34. Coen ES, Carpenter R, Martin C (1986) Transposable elements generate novel spatial patterns of gene expression in Antirrhinum majus. Cell 47: 285–296

35. Coen ES, Romero JM, Doyle S, Elliott R, Murphy G, Carpenter R (1990) floricaula: A homeotic gene required for flower development in Antirrhinum majus. Cell 63: 1311–1322

36. Coenen H, Viaene T, Vandenbussche M, Geuten K (2018) TM8 represses developmental timing in Nicotiana benthamiana and has functionally diversified in angiosperms. BMC Plant Biol 18: 129

37. Corley SB, Carpenter R, Copsey L, Coen E (2005) Floral asymmetry involves an interplay between TCP and MYB transcription factors in Antirrhinum. Proc Natl Acad Sci U S A 102: 5068–5073

38. Costa MMR, Fox S, Hanna AI, Baxter C, Coen E (2005) Evolution of regulatory interactions controlling floral asymmetry. Development 132: 5093–5101

39. Crawford BCW, Nath U, Carpenter R, Coen ES (2004) CINCINNATA controls both cell differentiation and growth in petal lobes and leaves of Antirrhinum. Plant Physiol 135: 244–53

40. Cubas P (2004) Floral zygomorphy, the recurring evolution of a successful trait. Bioessays 26: 1175–1184

41. Cullen E, Wang Q, Glover BJ (2023) How do you build a nectar spur? A transcriptomic comparison of nectar spur development in Linaria vulgaris and gibba development in Antirrhinum majus. Front Plant Sci 14: 1190373

42. Davies KM, Marshall GB, Bradley JM, Schwinn KE, Bloor SJ, Winefield CS, Martin CR (2006) Characterisation of aurone biosynthesis in Antirrhinum majus. Physiol Plant 128: 593–603

43. DeBono A, Yeats TH, Rose JKC, Bird D, Jetter R, Kunst L, Samuels L (2009) Arabidopsis LTPG Is a Glycosylphosphatidylinositol-Anchored Lipid Transfer Protein Required for Export of Lipids to the Plant Surface. Plant Cell 21: 1230–1238

44. Ding L, Yan S, Jiang L, Zhao W, Ning K, Zhao J, Liu X, Zhang J, Wang Q, Zhang X (2015) HANABA TARANU (HAN) Bridges Meristem and Organ Primordia Boundaries through PINHEAD, JAGGED, BLADE-ON-PETIOLE2 and CYTOKININ OXIDASE 3 during Flower Development in Arabidopsis. PLoS Genet 11: e1005479

45. Engelhorn J, Reimer JJ, Leuz I, Göbel U, Huettel B, Farrona S, Turck F (2012) Development-Related PcG target in the apex controls leaf margin architecture in Arabidopsis thaliana. Development 139: 2566–2575

46. Espinosa-Ruiz A, Martínez C, De Lucas M, Fàbregas N, Bosch N, Caño-Delgado AI, Prat S (2017) TOPLESS mediates brassinosteroid control of shoot boundaries and root meristem development in Arabidopsis thaliana. Development 144: 1619–1628

47. Galego L, Almeida J (2002) Role of DIVARICATA in the control of dorsoventral asymmetry in Antirrhinum flowers. Genes Dev 16: 880–891

48. Gampala SS, Kim TW, He JX, Tang W, Deng Z, Bai MY, Guan S, Lalonde S, Sun Y, Gendron JM, et al (2007) An essential role for 14-3-3 proteins in brassinosteroid signal transduction in Arabidopsis. Dev Cell 13: 177–189

49. Gaudin V, Lunness PA, Fobert PR, Towers M, Riou-Khamlichi C, Murray JAH, Coen E, Doonan JH (2000) The expression of D-cyclin genes defines distinct developmental zones in snapdragon apical meristems and is locally regulated by the Cycloidea gene. Plant Physiol 122: 1137

50. Gendron JM, Liu JS, Fan M, Bai MY, Wenkel S, Springer PS, Barton MK, Wang ZY (2012) Brassinosteroids regulate organ boundary formation in the shoot apical meristem of Arabidopsis. Proc Natl Acad Sci U S A 109: 21152–21157

51. Glover BJ, Perez-Rodriguez M, Martin C (1998) Development of several epidermal cell types can be specified by the same MYB-related plant transcription factor. Development 125: 3497–508

52. Gomez MD, Barro-Trastoy D, Escoms E, Saura-Sańchez M, Sańchez I, Briones-Moreno A, Vera-Sirera F, Carrera E, Ripoll JJ, Yanofsky MF, et al (2018) Gibberellins negatively modulate ovule number in plants. Development 145: dev163865

53. Gonçalves B, Hasson A, Belcram K, Cortizo M, Morin H, Nikovics K, Vialette-Guiraud A, Takeda S, Aida M, Laufs P, et al (2015) A conserved role for CUP-SHAPED COTYLEDON genes during ovule development. The Plant Journal 83: 732–742

54. Götz S, García-Gómez JM, Terol J, Williams TD, Nagaraj SH, Nueda MJ, Robles M, Talón M, Dopazo J, Conesa A (2008) High-throughput functional annotation and data mining with the Blast2GO suite. Nucleic Acids Res 36: 3420–3450

55. Grant CE, Bailey TL, Noble WS (2011) FIMO: Scanning for occurrences of a given motif. Bioinformatics 27: 1017–1018

56. Green AA, Kennaway JR, Hanna AI, Andrew Bangham J, Coen E (2010) Genetic control of organ shape and tissue polarity. PLoS Biol 8: e1000537

57. Han X, Hyun TK, Zhang M, Kumar R, Koh E ji, Kang BH, Lucas WJ, Kim JY (2014) Auxin-callose-mediated plasmodesmal gating is essential for tropic auxin gradient formation and signaling. Dev Cell 28: 132–146

58. Hao YJ, Song QX, Chen HW, Zou HF, Wei W, Kang XS, Ma B, Zhang WK, Zhang JS, Chen SY (2010) Plant NAC-type transcription factor proteins contain a NARD domain for repression of transcriptional activation. Planta 232: 1033–1043

59. Hasson A, Plessis A, Blein T, Adroher B, Grigg S, Tsiantis M, Boudaoud A, Damerval C, Laufs P (2011) Evolution and diverse roles of the CUP-SHAPED COTYLEDON genes in Arabidopsis leaf development. Plant Cell 23: 54–68

60. He JX, Gendron JM, Sun Y, Gampala SSL, Gendron N, Sun CQ, Wang ZY (2005) BZR1 is a transcriptional repressor with dual roles in brassinosteroid homeostasis and growth responses. Science 307: 1634–1638

61. Hendelman A, Stav R, Zemach H, Arazi T (2013) The tomato NAC transcription factor SlNAM2 is involved in flower-boundary morphogenesis. J Exp Bot 64: 5497–5507

62. Heo JB, Sung S, Assmann SM (2012) Ca 2+-dependent GTPase, extra-large G protein 2 (XLG2), promotes activation of DNA-binding protein related to vernalization 1 (RTV1), leading to activation of floral integrator genes and early flowering in Arabidopsis. Journal of Biological Chemistry 287: 8242–8253

63. Hibara KI, Karim MR, Takada S, Taoka KI, Furutani M, Aida M, Tasaka M (2006) Arabidopsis CUP-SHAPED COTYLEDON3 regulates postembryonic shoot meristem and organ boundary formation. Plant Cell 18: 2946–2957

64. Hileman LC (2014) Trends in flower symmetry evolution revealed through phylogenetic and developmental genetic advances. Philosophical Transactions of the Royal Society Series B, Biological Sciences 369: 20130348

65. Huang G, Han M, Jian L, Chen Y, Sun S, Wang X, Wang Y (2020) An ETHYLENE INSENSITIVE3-LIKE1 protein directly targets the GEG promoter and mediates ethylene-induced ray petal elongation in Gerbera hybrida. Front Plant Sci 10: 1737

66. Ichimura K, Niki T, Matoh M, Nakayama M (2021) High temperature under low light conditions suppresses anthocyanin biosynthesis in snapdragon petals associated with decreased sugar levels. Sci Hortic 290: 110510

67. Israeli A, Reed JW, Ori N (2020) Genetic dissection of the auxin response network. Nat Plants 6: 1082–1090

68. Jackson D (1992) In situ hybridization in plants. Mol Plant Pathol. Oxford Academic, Oxford, pp 163–174

69. Jackson D, Roberts K, Martin C (1992) Temporal and spatial control of expression of anthocyanin biosynthetic genes in developing flowers of Antirrhinum majus. The Plant Journal 2: 425–434

70. Jaffé FW, Tattersall A, Glover BJ (2007) A truncated MYB transcription factor from Antirrhinum majus regulates epidermal cell outgrowth. J Exp Bot 58: 1515–1524

71. Javelle M, Timmermans MCP (2012) In situ localization of small RNAs in plants by using LNA probes. Nat Protoc 7: 533–541

72. Jiang R, Yuan W, Yao W, Jin X, Wang X, Wang Y (2022) A regulatory GhBPE-GhPRGL module maintains ray petal length in Gerbera hybrida. Molecular Horticulture 2: 9

73. Jiao K, Li X, Guo W, Su S, Da Luo (2017) High-throughput RNA-seq data analysis of the single nucleotide polymorphisms (SNPs) and zygomorphic flower development in pea (Pisum sativum L.). Int J Mol Sci 18: 2710

74. Kamiuchi Y, Yamamoto K, Furutani M, Tasaka M, Aida M (2014) The CUC1 and CUC2 genes promote carpel margin meristem formation during Arabidopsis gynoecium development. Front Plant Sci 5: 165

75. Kawamura E, Horiguchi G, Tsukaya H (2010) Mechanisms of leaf tooth formation in Arabidopsis. The Plant Journal 62: 429–441

76. Kim D, Langmead B, Salzberg SL (2015) HISAT: a fast spliced aligner with low memory requirements. Nat Methods 12: 357–360

77. Kirchhelle C, Chow CM, Foucart C, Neto H, Stierhof YD, Kalde M, Walton C, Fricker M, Smith RS, Jérusalem A, et al (2016) The specification of geometric edges by a plant Rab GTPase is an essential cell-patterning principle during organogenesis in Arabidopsis. Dev Cell 36: 386–400

78. Krueger F (2023) TrimGalore. V0.6.7, https://github.com/FelixKrueger/TrimGalore

79. Kurakawa T, Ueda N, Maekawa M, Kobayashi K, Kojima M, Nagato Y, Sakakibara H, Kyozuka J (2007) Direct control of shoot meristem activity by a cytokinin-activating enzyme. Nature 445: 652–655

80. Kuroha T, Tokunaga H, Kojima M, Ueda N, Ishida T, Nagawa S, Fukuda H, Sugimoto K, Sakakibara H (2009) Functional analyses of LONELY GUY cytokinin-activating enzymes reveal the importance of the direct activation pathway in Arabidopsis. Plant Cell 21: 3152–3169

81. Laufs P, Peaucelle A, Morin H, Traas J (2004) MicroRNA regulation of the CUC genes is required for boundary size control in Arabidopsis meristems. Development 131: 4311–4322

82. Leonte G (2020) Characterization of cluster-III HIPP proteins targeted to plasmodesmata in Arabidopsis thaliana. Doctoral dissertation. Freie Universität Berlin

83. Levy YY, Mesnage S, Mylne JS, Gendall AR, Dean C (2002) Multiple roles of Arabidopsis VRN1 in vernalization and flowering time control. Science 297: 243

84. Li C, Yu W, Xu J, Lu X, Liu Y (2022) Anthocyanin Biosynthesis Induced by MYB Transcription Factors in Plants. Int J Mol Sci 23: 11701

85. Li L, Zhang W, Li LZN, Peng J, Wang Y, Zhong C, Yang Y, Sun S, Liang S, Wang X (2015) Transcriptomic insights into antagonistic effects of gibberellin and abscisic acid on petal growth in Gerbera hybrida. Front Plant Sci 6: 168

86. Li M (2002) Molecular and Genetic Characterization of New MADS-box Genes in Antirrhinum majus. Doctoral thesis. Universität zu Köln

87. Li M, Zhang D, Gao Q, Luo Y, Zhang H, Ma B, Chen C, Whibley A, Zhang Y, Cao Y, et al (2019) Genome structure and evolution of Antirrhinum majus L. Nat Plants 5: 174–183

88. Love MI, Huber W, Anders S (2014) Moderated estimation of fold change and dispersion for RNA-seq data with DESeq2. Genome Biol 15: 550

89. Luo D, Carpenter R, Copsey L, Vincent C, Clark J, Coen E (1999) Control of organ asymmetry in flowers of Antirrhinum. Cell 99: 367–376

90. Luo D, Carpenter R, Vincent C, Copsey L, Coen E (1996) Origin of floral asymmetry in Antirrhinum. Nature 383: 794–799

91. Luo J, Zhou JJ, Zhang JZ (2018) Aux/IAA gene family in plants: Molecular structure, regulation, and function. Int J Mol Sci 19: 259

92. Ma N, Xue J, Li Y, Liu X, Dai F, Jia W, Luo Y, Gao J (2008) Rh-PIP2;1, a rose aquaporin gene, is involved in ethylene-regulated petal expansion. Plant Physiol 148: 894–907

93. Mallory AC, Dugas D V., Bartel DP, Bartel B (2004) MicroRNA regulation of NAC-domain targets is required for proper formation and separation of adjacent embryonic, vegetative, and floral organs. Curr Biol 14: 1035–1046

94. Manrique S, Caselli F, Matías-Hernández L, Franks RG, Colombo L, Gregis V (2023) Assessing the role of REM13, REM34 and REM46 during the transition to the reproductive phase in Arabidopsis thaliana. Plant Mol Biol 112: 179–193

95. Martin C, Prescott A, Mackay S, Bartlett J, Vrijlandt E (1991) Control of anthocyanin biosynthesis in flowers of Antirrhinum majus. The Plant Journal 1: 37–49

96. Matias-Hernandez L, Battaglia R, Galbiati F, Rubes M, Eichenberger C, Grossniklaus U, Kater MM, Colombo L (2010) VERDANDI is a direct target of the MADS domain ovule identity complex and affects embryo sac differentiation in Arabidopsis. Plant Cell 22: 1702–1715

97. Mendes MA, Guerra RF, Castelnovo B, Silva-Velazquez Y, Morandini P, Manrique S, Baumann N, Groß-Hardt R, Dickinson H, Colombo L (2016) Live and let die: A REM complex promotes fertilization through synergid cell death in Arabidopsis. Development 143: 2780–2790

98. Nath U, Crawford BCW, Carpenter R, Coen E (2003) Genetic control of surface curvature. Science 299: 1404–1407

99. Nicolas A, Maugarny-Calès A, Adroher B, Chelysheva L, Li Y, Burguet J, Bågman AM, Smit ME, Brady SM, Li Y, et al (2022) De novo stem cell establishment in meristems requires repression of organ boundary cell fate. Plant Cell 34: 4738–4759

100. Nikovics K, Blein T, Peaucelle A, Ishida T, Morin H, Aida M, Laufs P (2006) The balance between the MIR164A and CUC2 genes controls leaf margin serration in Arabidopsis. Plant Cell 18: 2929–2945

101. Ono E, Fukuchi-Mizutani M, Nakamura N, Fukui Y, Yonekura-Sakakibara K, Yamaguchi M, Nakayama T, Tanaka T, Kusumi T, Tanaka Y (2006) Yellow flowers generated by expression of the aurone biosynthetic pathway. Proc Natl Acad Sci U S A 103: 11075–11080

102. Pan ZJ, Nien YC, Shih YA, Chen TY, Lin WD, Kuo WH, Hsu HC, Tu SL, Chen JC, Wang CN (2022) Transcriptomic analysis suggests auxin regulation in dorsal–ventral petal asymmetry of wild progenitor Sinningia speciosa. Int J Mol Sci 23: 2073

103. Pei H, Ma N, Tian J, Luo J, Chen J, Li J, Zheng Y, Chen X, Fei Z, Gao J (2013) An NAC transcription factor controls ethylene-regulated cell expansion in flower petals. Plant Physiol 163: 775–791

104. Perez-Rodriguez M, Jaffe FW, Butelli E, Glover BJ, Martin C (2005) Development of three different cell types is associated with the activity of a specific MYB transcription factor in the ventral petal of Antirrhinum majus flowers. Development 132: 359–370

105. Petroni K, Tonelli C (2011) Recent advances on the regulation of anthocyanin synthesis in reproductive organs. Plant Science 181: 219–229

106. Preston JC, Hileman LC (2009) Developmental genetics of floral symmetry evolution. Trends Plant Sci 14: 147–154

107. Preston JC, Hileman LC, Cubas P (2011) Reduce, reuse, and recycle: developmental evolution of trait diversification. Am J Bot 98: 397–403

108. Raimundo J, Sobral R, Bailey P, Azevedo H, Galego L, Almeida J, Coen E, Costa MMR (2013) A subcellular tug of war involving three MYB-like proteins underlies a molecular antagonism in Antirrhinum flower asymmetry. Plant J 75: 527–538

109. Raman S, Greb T, Peaucelle A, Blein T, Laufs P, Theres K (2008) Interplay of miR164, CUP-SHAPED COTYLEDON genes and LATERAL SUPPRESSOR controls axillary meristem formation in Arabidopsis thaliana. The Plant Journal 55: 65–76

110. Rebocho AB, Kennaway JR, Bangham JA, Coen E (2017a) Formation and Shaping of the Antirrhinum Flower through Modulation of the CUP Boundary Gene. Current Biology 27: 2610–2622.e3

111. Rebocho AB, Southam P, Kennaway RJ, Bangham AJ, Coen E (2017b) Generation of shape complexity through tissue conflict resolution. Elife 6: e20156

112. Richter R, Kinoshita A, Vincent C, Martinez-Gallegos R, Gao H, van Driel AD, Hyun Y, Mateos JL, Coupland G (2019) Floral regulators FLC and SOC1 directly regulate expression of the B3-type transcription factor TARGET of FLC and SVP 1 at the Arabidopsis shoot apex via antagonistic chromatin modifications. PLoS Genet 15: e1008065

113. Riou-Khamlichi C, Huntley R, Jacqmard A, Murray JAH (1999) Cytokinin activation of Arabidopsis cell division through a D-type cyclin. Science 283: 1541–1544

114. Rodas AL, Roque E, Hamza R, Gómez-Mena C, Minguet EG, Wen J, Mysore KS, Beltrán JP, Cañas LA (2021) MtSUPERMAN plays a key role in compound inflorescence and flower development in Medicago truncatula. The Plant Journal 105: 816–830

115. Romanel EAC, Schrago CG, Couñago RM, Russo CAM, Alves-Ferreira M (2009) Evolution of the B3 DNA binding superfamily: new insights into REM family gene diversification. PLoS One 4: e5791

116. Rosin FM, Kramer EM (2009) Old dogs, new tricks: Regulatory evolution in conserved genetic modules leads to novel morphologies in plants. Dev Biol 332: 25–35

117. Sakai H, Medrano LJ, Meyerowitz EM (1995) Role of SUPERMAN in maintaining Arabidopsis floral whorl boundaries. Nature 378: 199–203

118. Salehin M, Bagchi R, Estelle M (2015) ScfTIR1/AFB-based auxin perception: Mechanism and role in plant growth and development. Plant Cell 27: 9–19

119. Schwinn K, Venail J, Shang Y, Mackay S, Alm V, Butelli E, Oyama R, Bailey P, Davies K, Martin C (2006) A small family of MYB-regulatory genes controls floral pigmentation intensity and patterning in the genus Antirrhinum. Plant Cell 18: 831–851

120. Shao J, Meng J, Wang F, Shou B, Chen Y, Xue H, Zhao J, Qi Y, An L, Yu F, et al (2020) NGATHA-LIKEs control leaf margin development by repressing CUP-SHAPED COTYLEDON2 transcription. Plant Physiol 184: 345–358

121. Sommer H, Saedler H (1986) Structure of the chalcone synthase gene of Antirrhinum majus. Molecular and General Genetics 202: 429–434

122. Souer E, Van Houwelingen A, Kloos D, Mol J, Koes R (1996) The no apical meristem gene of Petunia is required for pattern formation in embryos and flowers and is expressed at meristem and primordia boundaries. Cell 85: 159–170

123. Suzuki H, Nakayama T, Yonekura-Sakakibara K, Fukui Y, Nakamura N, Nakao M, Tanaka Y, Yamaguchill MA, Kusumi T, Nishino T (2001) Malonyl-CoA:anthocyanin 5-O-glucoside-6‴-O-malonyltransferase from scarlet sage (Salvia splendens) flowers. Enzyme purification, gene cloning, expression, and characterization. Journal of Biological Chemistry 276: 49013–49029

124. Suzuki H, Sawada S, Watanabe K, Nagae S, Yamaguchi MA, Nakayama T, Nishino T (2004) Identification and characterization of a novel anthocyanin malonyltransferase from scarlet sage (Salvia splendens) flowers: an enzyme that is phylogenetically separated from other anthocyanin acyltransferases. Plant Journal 38: 994–1003

125. Swaminathan K, Peterson K, Jack T (2008) The plant B3 superfamily. Trends Plant Sci 13: 647–655

126. Thiele K, Wanner G, Kindzierski V, Jürgens G, Mayer U, Pachl F, Assaad FF (2009) The timely deposition of callose is essential for cytokinesis in Arabidopsis. Plant Journal 58: 13–26

127. Tokunaga H, Kojima M, Kuroha T, Ishida T, Sugimoto K, Kiba T, Sakakibara H (2012) Arabidopsis lonely guy (LOG) multiple mutants reveal a central role of the LOG-dependent pathway in cytokinin activation. The Plant Journal 69: 355–365

128. Tsuwamoto R, Fukuoka H, Takahata Y (2008) GASSHO1 and GASSHO2 encoding a putative leucine-rich repeat transmembrane-type receptor kinase are essential for the normal development of the epidermal surface in Arabidopsis embryos. Plant Journal 54: 30–42

129. Valoroso MC, Lucibelli F, Aceto S (2022) Orchid NAC transcription factors: a focused analysis of CUPULIFORMIS genes. Genes (Basel) 13: 2293

130. Vialette-Guiraud ACM, Adam H, Finet C, Jasinski S, Jouannic S, Scutt CP (2011) Insights from ANA-grade angiosperms into the early evolution of CUP-SHAPED COTYLEDON genes. Ann Bot 107: 1511–1519

131. Vialette-Guiraud ACM, Chauvet A, Gutierrez-Mazariegos J, Eschstruth A, Ratet P, Scutt CP (2016) A conserved role for the NAM/miR164 developmental module reveals a common mechanism underlying carpel margin fusion in monocarpous and syncarpous eurosids. Front Plant Sci 6: 1239

132. Vincent CA, Coen ES (2004) A temporal and morphological framework for flower development in Antirrhinum majus. Botany 82: 681–690

133. Wang J, Bao J, Zhou B, Li M, Li X, Jin J (2021) The osa-miR164 target OsCUC1 functions redundantly with OsCUC3 in controlling rice meristem/organ boundary specification. New Phytologist 229: 1566–1581

134. Wang X, Shen C, Meng P, Tan G, Lv L (2021) Analysis and review of trichomes in plants. BMC Plant Biol 21: 70

135. Weir I, Lu J, Cook H, Causier B, Schwarz-Sommer Z, Davies B (2004) CUPULIFORMIS establishes lateral organ boundaries in Antirrhinum. Development 131: 915–922

136. Wessinger CA, Hileman LC (2020) Parallelism in flower evolution and development. 51: 387–408

137. Xing S, Rosso MG, Zachgo S (2005) ROXY1, a member of the plant glutaredoxin family, is required for petal development in Arabidopsis thaliana. Development 132: 1555–1565

138. Xing S, Zachgo S (2008) ROXY1 and ROXY2, two Arabidopsis glutaredoxin genes, are required for anther development. Plant Journal 53: 790–801

139. Xu Y, Prunet N, Gan E-S, Wang Y, Stewart D, Wellmer F, Huang J, Yamaguchi N, Tatsumi Y, Kojima M, et al (2018) SUPERMAN regulates floral whorl boundaries through control of auxin biosynthesis. EMBO J 37: e97499

140. Yamaguchi N, Huang J, Tatsumi Y, Abe M, Sugano SS, Kojima M, Takebayashi Y, Kiba T, Yokoyama R, Nishitani K, et al (2018) Chromatin-mediated feed-forward auxin biosynthesis in floral meristem determinacy. Nat Commun 9: 5290

141. Yamaguchi N, Huang J, Xu Y, Tanoi K, Ito T (2017) Fine-tuning of auxin homeostasis governs the transition from floral stem cell maintenance to gynoecium formation. Nat Commun 8: 1125

142. Yan H, Pei X, Zhang H, Li X, Zhang X, Zhao M, Chiang VL, Sederoff RR, Zhao X (2021) Myb-mediated regulation of anthocyanin biosynthesis. Int J Mol Sci 22: 3103

143. Yang W, Cortijo S, Korsbo N, Roszak P, Schiessl K, Gurzadyan A, Wightman R, Jönsson H, Meyerowitz E (2021) Molecular mechanism of cytokinin-activated cell division in Arabidopsis. Science 371: 1350–1355

144. Yu Y, Qiao L, Chen J, Rong Y, Zhao Y, Cui X, Xu J, Hou X, Dong CH (2020) Arabidopsis REM16 acts as a B3 domain transcription factor to promote flowering time via directly binding to the promoters of SOC1 and FT. Plant Journal 103: 1386–1398

145. Zheng G, Wei W, Li Y, Kan L, Wang F, Zhang X, Li F, Liu Z, Kang C (2019) Conserved and novel roles of miR164-CUC2 regulatory module in specifying leaf and floral organ morphology in strawberry. New Phytologist 224: 480–492

146. Zhong J, Powell S, Preston JC (2016) Organ boundary NAC-domain transcription factors are implicated in the evolution of petal fusion. Plant Biol 18: 893–902

147. Zschiesche W, Barth O, Daniel K, Böhme S, Rausche J, Humbeck K (2015) The zinc-binding nuclear protein HIPP3 acts as an upstream regulator of the salicylate-dependent plant immunity pathway and of flowering time in Arabidopsis thaliana. New Phytologist 207: 1084

