## Supplementary Figures and Tables for "Antirrhinum flower shape: unravelling gene expression across developmental axes and boundaries"

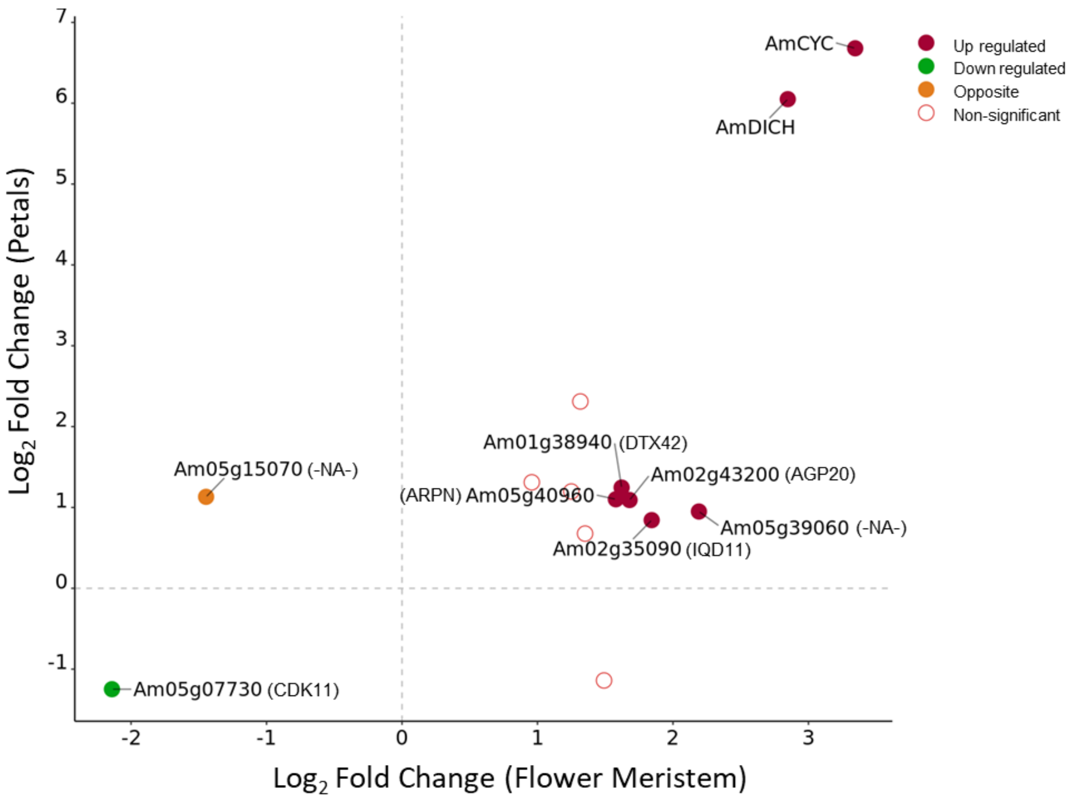


**Supplementary Figure 1.** Correlation between differentially expressed genes (DEGs) obtained from two flower developmental stages. DEGs were obtained from the comparison between dorsal and ventral regions of the flower meristem and from the comparison between dorsal and ventral petals. All DEGs with FDR<<0.05 in common between the two stages were plotted. Filled circles represent significant genes, FDR<0.01, with circles filled in red representing dorsal genes (up regulated in both analysis), green filled circles representing ventral genes (down regulated in both analysis) and an orange filled circle representing a gene that has opposite expression in between analysis. The genes identified included *AmCYC (Am08g22680)* and *AmDICH (Am06g32830)*, while the remaining genes were labelled with the abbreviated gene description of their homologues obtained from a BLAST search. Genes without BLAST results for homologues were labelled as "-NA-".

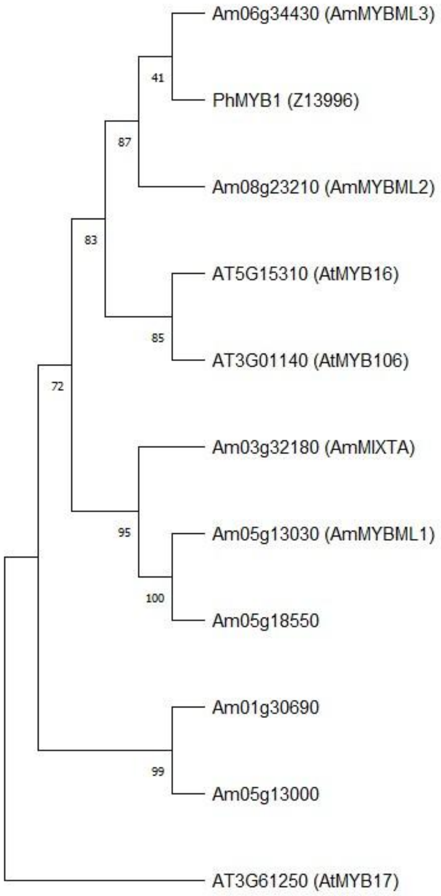


**Supplementary Figure 2.** Phylogenetic analysis of MYBML proteins. AmMYBL1 sequence was used in a blast search against the translated proteins from the *Antirrhinum majus* genome. Homologues of AmMYBML1 were obtained for *Arabidopsis thaliana* by performing a blast in TAIR BLAST 2.9.0+ and from a previous analysis done by Perez-Rodriguez et al. (2005). The closest BLAST homologues of Am05g18550 and Am01g30690 were MYB160 and MYB16 from *Arabidopsis*, respectively. Protein sequences of the MYBML family from *Antirrhinum majus* (Am), *Arabidopsis thaliana* (At) and *Petunia hybrida* (Ph) were aligned using MUSCLE in MEGA11. A phylogenetic tree was inferred using the Maximum Likelihood method and JTT matrix-based model with 100 bootstraps.

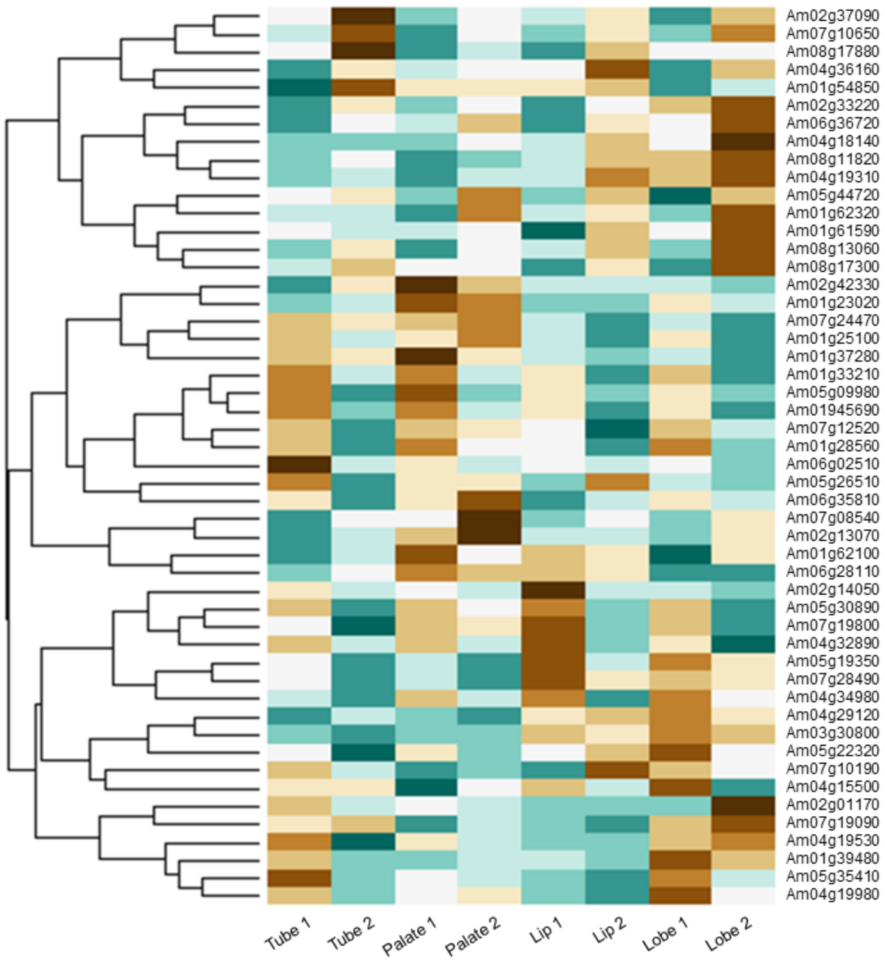


**Supplementary Figure 3.** Heatmap genes expressed in ventral petal regions used as control. Fifty genes were randomly selected from a pool of genes with expression in all regions of the ventral petal, tube, palate, lip and lobe. Normalised expression of two biological replicates was retrieved and plotted in a heatmap with hierarchical clustering. The randomly chosen genes showed an absence of clustering according to the ventral petal regions.

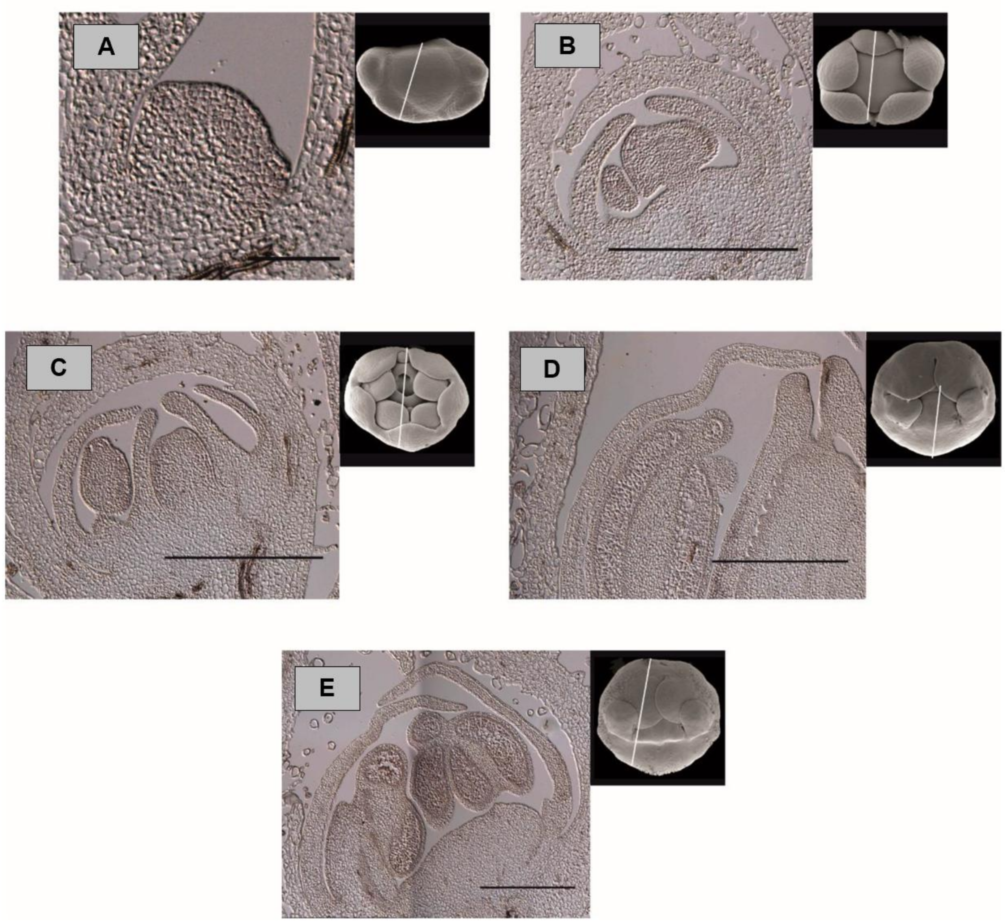


**Supplementary Figure 4.** Several developmental stages of *Antirrhinum majus* flower meristems at (A) 6 DAI; (B) 7 DAI; (C) 10 DAI; (D) 12 DAI; (E) 14 DAI hybridized with a representative sense probe. The ventral petal is positioned to the left and dorsal petal to the right. The diagram at the right of each image shows a picture of the flower bud at the corresponding developmental stage and the section plane of the respective image. Scale bars correspond to 500 μm, except for A, where it corresponds to 100 μm.

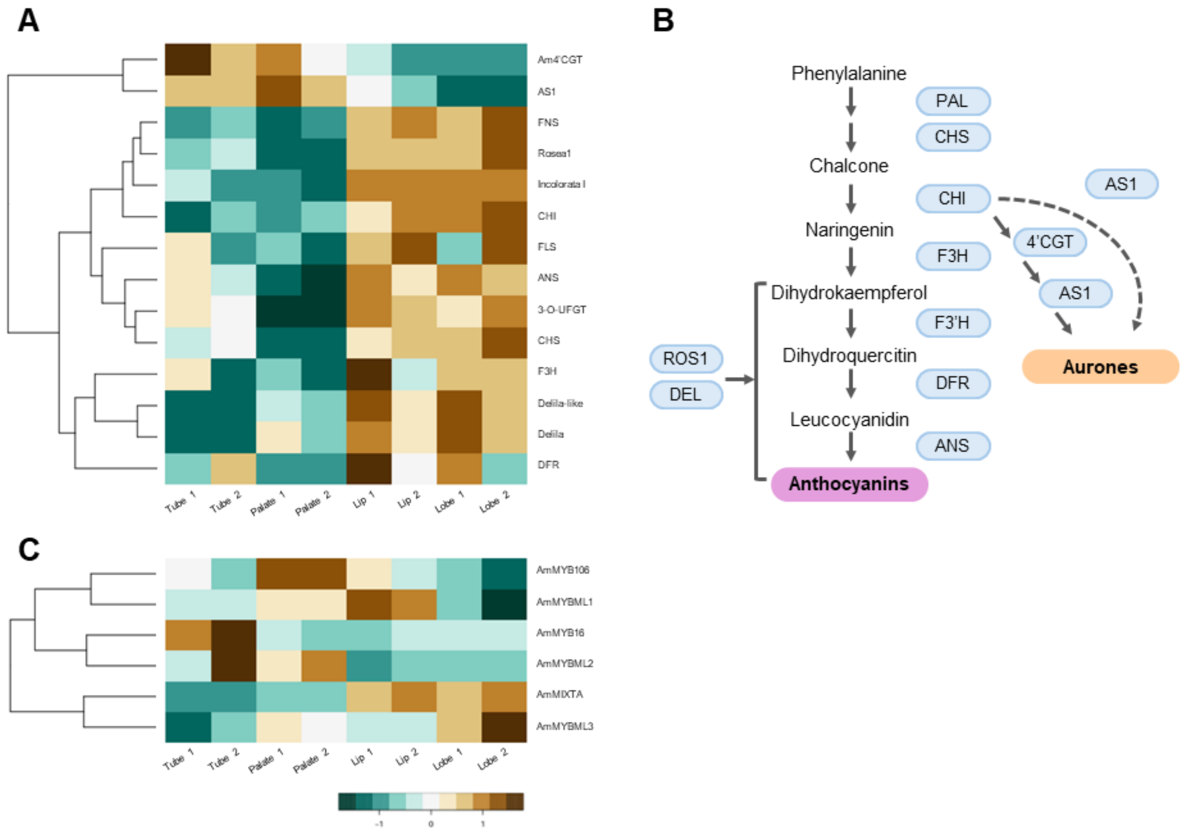


**Supplementary Figure 5.** Expression of genes related to colour and production of specific cell types in the ventral petal. **(A)** Heatmap with hierarchical clustering of genes of involved in the biosynthesis of anthocyanins and aurones across samples of tube, palate, lip, and lobe. Normalised expression of two biological replicates is shown. **(B)** Scheme of the biosynthetic pathway that produces anthocyanins and aurones in *Antirrhinum* (Martin et al., 1991; Jackson et al., 1992; Davies et al., 2006; Petroni and Tonelli, 2011). **(C)** The normalised expression of genes from the *MYBML* family was analysed in expression in different parts of ventral petal.

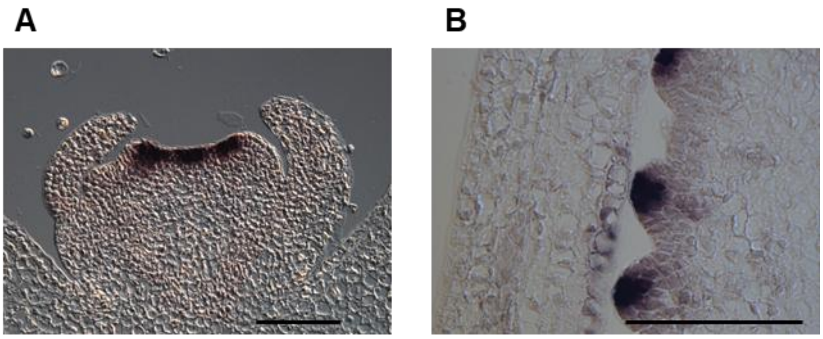

**Supplementary Figure 6.** Expression pattern of *LOG5* detected by *in situ* hybridisation. **(A)** At 7 days after meristem initiation (DAI) *LOG5* is expressed in the organ primordia of stamens and carpels. **(B)** At 14 DAI, *LOG5* is expressed in ovules and absent from ovule boundaries. Scale 100 μM.

### Supplementary Tables

**Supplementary Table 1.** Expanded data on samples of different tissues and stages used in transcriptomic analysis. Samples from the flower meristem (FM) at 5 days after meristem initiation (DAI) and different regions of the ventral petal (Tube, Palate, Lip, Lobe) were isolated by laser microdissection (LMD) from a *cyc dich* mutant. The different petals of WT flowers (dorsal, lateral and ventral) were dissected of 5 mm wide buds, whereas whole corollas were used for the *cyc dich* and *cyc dich div* mutant samples. RNA samples were sent for sequencing varying in read size (50 or 150 base pairs) and library type (Single or Paired). The number of reads after trimming are shown in millions (M) of reads. Mapping was done against *Antirrhinum majus* genome, and the percentage of reads mapped are shown.

| Sample name | Stage | Sample type | Read size | Library type | Trimmed reads (M) | Mapped % |
|---|---|---|---|---|---|---|
| Dorsal_FM_1 | 5 DAI | LMD tissue | 150 | Paired | 21.2 | 77.1 |
| Dorsal_FM_2 | 5 DAI | LMD tissue | 150 | Paired | 12.1 | 85.1 |
| Ventral_FM_1 | 5 DAI | LMD tissue | 150 | Paired | 22 | 85.1 |
| Ventral_FM_2 | 5 DAI | LMD tissue | 150 | Paired | 12.4 | 72.7 |
| Dorsal_P_1 | 5 mm wide buds | Petal | 50 | Single | 31.6 | 98.3 |
| Dorsal_P_2 | 5 mm wide buds | Petal | 50 | Single | 18.8 | 98.3 |
| Dorsal_P_3 | 5 mm wide buds | Petal | 50 | Single | 21.0 | 98.2 |
| Lateral_P_1 | 5 mm wide buds | Petal | 50 | Single | 30.0 | 98.3 |
| Lateral_P_2 | 5 mm wide buds | Petal | 50 | Single | 22.5 | 98.4 |
| Lateral_P_3 | 5 mm wide buds | Petal | 50 | Single | 29.7 | 98.2 |
| Ventral_P_1 | 5 mm wide buds | Petal | 50 | Single | 25.3 | 98.2 |
| Ventral_P_2 | 5 mm wide buds | Petal | 50 | Single | 45.4 | 98.3 |
| Ventral_P_3 | 5 mm wide buds | Petal | 50 | Single | 18.8 | 98.1 |
| cycdich_1 | 5 mm wide buds | Corolla | 50 | Single | 28.0 | 98.1 |
| cycdich_2 | 5 mm wide buds | Corolla | 50 | Single | 28.0 | 98.1 |
| cycdich_3 | 5 mm wide buds | Corolla | 50 | Single | 32.9 | 98.0 |
| cycdichdiv_1 | 5 mm wide buds | Corolla | 50 | Single | 23.9 | 93.8 |
| cycdichdiv_2 | 5 mm wide buds | Corolla | 50 | Single | 15.8 | 90.4 |
| cycdichdiv_3 | 5 mm wide buds | Corolla | 50 | Single | 25.0 | 94.2 |
| Tube_1 | 6.5 mm wide buds | LMD tissue | 50 | Single | 35.4 | 98.5 |
| Tube_2 | 6.5 mm wide buds | LMD tissue | 50 | Single | 21.6 | 98.3 |
| Palate_1 | 6.5 mm wide buds | LMD tissue | 50 | Single | 21.4 | 98.4 |
| Palate_2 | 6.5 mm wide buds | LMD tissue | 50 | Single | 32.3 | 98.2 |
| Lip_1 | 6.5 mm wide buds | LMD tissue | 50 | Single | 9.50 | 98.3 |
| Lip_2 | 6.5 mm wide buds | LMD tissue | 50 | Single | 44.9 | 98.4 |
| Lobe_1 | 6.5 mm wide buds | LMD tissue | 50 | Single | 19.3 | 91.2 |
| Lobe_2 | 6.5 mm wide buds | LMD tissue | 50 | Single | 36.3 | 98.5 |
| WT_1 | 3.5 mm wide buds | Bud | 50 | Single | 13.8 | 90.0 |
| WT_2 | 3.5 mm wide buds | Bud | 50 | Single | 34.2 | 90.3 |
| cup_1 | 3.5 mm wide buds | Bud | 50 | Single | 58.4 | 90.2 |
| cup_2 | 3.5 mm wide buds | Bud | 50 | Single | 13.7 | 89.9 |

**Supplementary Table: 2.** AmCYC binding sites in promoters of genes more expressed in dorsal regions of the flower meristem. Promoter sequences were defined as 1000 base pairs before the initiation codon and analysed for AmCYC binding sites (5'-GGNCCCNC-3') (Costa et al., 2005) using FIMO. Genes without BLAST results for homologues were labelled as "-NA-".

| Gene Description | GeneID | Strand | Start | End | p-value | q-value | Matched Sequence |
|---|---|---|---|---|---|---|---|
| *DTX42* | *Am01g38940* | - | 357 | 364 | 0.000131 | 0.445 | GGTCCCAC |
| *TSJT1* | *Am01g62890* | - | 806 | 813 | 9.17e-05 | 0.352 | GGACCCGC |
| | | - | 868 | 875 | 0.000131 | 0.445 | GGTCCCAC |
| *AGP20* | *Am02g43200* | - | 825 | 832 | 9.17e-05 | 0.352 | GGACCCCC |
| *GSL4* | *Am03g05080* | + | 716 | 723 | 2.67e-05 | 0.192 | GGGCCCCC |
| | | - | 714 | 721 | 2.67e-05 | 0.192 | GGGCCCCC |
| | | + | 727 | 734 | 2.67e-05 | 0.192 | GGCCCCCC |
| *MYB60* | *Am03g22020* | + | 498 | 505 | 2.67e-05 | 0.192 | GGCCCCCC |
| *ADH2* | *Am05g05520* | + | 632 | 639 | 9.17e-05 | 0.352 | GGGCCCAC |
| | | - | 630 | 637 | 9.17e-05 | 0.352 | GGGCCCTC |
| *-NA-* | *Am05g39060* | - | 149 | 156 | 9.17e-05 | 0.352 | GGCCCCTC |
| *PP2C24* | *Am06g27840* | + | 570 | 577 | 9.17e-05 | 0.352 | GGCCCCAC |
| *ROXY2* | *Am06g39760* | - | 928 | 935 | 9.17e-05 | 0.352 | GGCCCCTC |
| *-NA-* | *Am08g07490* | - | 249 | 256 | 2.67e-05 | 0.192 | GGCCCCCC |
| | | - | 250 | 257 | 2.67e-05 | 0.192 | GGGCCCCC |
| *AmCYC* | *Am08g22680* | - | 541 | 548 | 2.67e-05 | 0.192 | GGCCCCCC |
| | | - | 542 | 549 | 2.67e-05 | 0.192 | GGGCCCCC |

**Supplementary Table: 3.** DIV-binding sites in promoters of genes possibly regulated by AmDIV. Genes possibly regulated by AmDIV were defined as being more expressed the ventral petal and *cyc dich* mutant (WT dorsal vs. ventral and WT dorsal vs. *cyc dich,* respectively) and down regulated in *cyc dich div (cyc dich* vs. *cyc dich div)*. Promoter sequences were defined as 1000 base pairs before the initiation codon and analysed for AmDIV binding sites (5'- GATAAG -3') (Raimundo et al., 2013) using FIMO. Genes without BLAST results for homologues were labelled as "-NA-".

| Gene Description | GeneID | Strand | Start | End | p-value | q-value | Matched Sequence |
|---|---|---|---|---|---|---|---|
| *--NA--* | *Am01g29280* | + | 378 | 383 | 0.000289 | 0.787 | GATAAG |
| *MYB16* | *Am01g30690* | - | 474 | 479 | 0.000289 | 0.787 | GATAAG |
| | *Am01g30690* | + | 547 | 552 | 0.000289 | 0.787 | GATAAG |
| *JGB* | *Am01g31630* | - | 158 | 163 | 0.000289 | 0.787 | GATAAG |
| *--NA--* | *Am02g43010* | + | 821 | 826 | 0.000289 | 0.787 | GATAAG |
| *GATA9* | *Am02g44520* | - | 136 | 141 | 0.000289 | 0.787 | GATAAG |
| *AAE* | *Am03g10330* | + | 485 | 490 | 0.000289 | 0.787 | GATAAG |
| *HIPP22* | *Am04g17710* | - | 8 | 13 | 0.000289 | 0.787 | GATAAG |
| | *Am04g17710* | + | 340 | 345 | 0.000289 | 0.787 | GATAAG |
| *AAE* | *Am04g36320* | - | 112 | 117 | 0.000289 | 0.787 | GATAAG |
| *5MAT2* | *Am04g36910* | - | 862 | 867 | 0.000289 | 0.787 | GATAAG |
| *bHLH126* | *Am07g19180* | + | 183 | 188 | 0.000289 | 0.787 | GATAAG |
| | *Am07g19180* | - | 829 | 834 | 0.000289 | 0.787 | GATAAG |
| *BBX20* | *Am08g04660* | - | 401 | 406 | 0.000289 | 0.787 | GATAAG |
| | *Am08g04660* | - | 474 | 479 | 0.000289 | 0.787 | GATAAG |
| *RADL6* | *Am08g05030* | + | 190 | 195 | 0.000289 | 0.787 | GATAAG |
| *CAHC* | *Am08g20160* | - | 62 | 67 | 0.000289 | 0.787 | GATAAG |

**Supplementary Table 4.** BZR1 binding sites in the *AmCUP* promoter. *AmCUP* promoter sequence was defined as 1000 base pairs before the initiation codon and analysed for BZR1 binding sites (5'- CGTGCG -3') from *Arabidopsis thaliana* (He et al., 2005) using FIMO.

| Motif | Sequence Name | Strand | Start | End | p-value | q-value | Matched Sequence |
|---|---|---|---|---|---|---|---|
| CGTGYG | Am03g10700 | - | 517 | 522 | 0.000355 | 0.141 | CGTGTG |
| | | + | 544 | 549 | 0.000355 | 0.141 | CGTGTG |
| | | - | 554 | 559 | 0.000355 | 0.141 | CGTGTG |
| | | - | 696 | 701 | 0.000355 | 0.141 | CGTGTG |
